# A Rare Multipotent Peg-like Epithelial Cell is a Candidate Cell-of-Origin for High-Grade Serous Ovarian Cancer

**DOI:** 10.1101/2025.09.23.677404

**Authors:** Megan L. Ritting, Wenmei Yang, Syed Mohammed Musheer Aalam, Hui Zhao, Liang Feng, Jianning Song, Mihai G. Dumbrava, Wazim M. Ismail, Dong-Gi Mun, Chunling Hu, Opalina Roy, Keyur Chaludiya, Geng Xian Shi, Daphne Norma Crasta, Kenneth Schaufelberger, Jeffrey R. Janus, Guruprasad Kalthur, S. John Weroha, Scott H. Kaufmann, Anguraj Sadanandam, David J. H. F. Knapp, Chen Wang, Akhilesh Pandey, Alexandre Gaspar-Maia, Fergus J. Couch, Mark E. Sherman, Jamie N. Bakkum-Gamez, Nagarajan Kannan

## Abstract

To illuminate the origins of high-grade serous ovarian cancer (HGSOC), the most lethal and common form of ovarian cancer, we have created a comprehensive living organoid biobank of human fallopian tube tissue, which is thought to be the origin of this cancer. Through optimized culture protocols and integrated multi-omic profiling—including single-cell RNA sequencing, chromatin accessibility (ATAC) analysis, proteomics, and secretomics—we assembled the largest molecular atlas of the fallopian tube epithelium to date. This resource revealed diverse epithelial lineages and regulatory networks, including a rare, multipotent epithelial subpopulation with hybrid epithelial–mesenchymal features. Spatially localized to the basal epithelium and resembling mesonephric developmental precursors, these cells exhibit transcriptomic and proteomic similarities to the mesenchyme-like subtype of HGSOC, implicating them as potential cells-of-origin. Their molecular identity is preserved in organoid models, enabling future mechanistic and translational studies. This resource, which advances fundamental understanding of epithelial hierarchy and cancer susceptibility, provides a platform to inform early detection and prevention strategies for aggressive forms of ovarian cancer.

**Highlights:**

- Establishment of a clinically annotated fallopian tube organoid biobank enables delineation of epithelial lineage hierarchies and differentiation capacity.
- Multi-omics integration defines robust, lineage-specific transcriptional and regulatory networks in the fallopian tube epithelium.
- A rare basal epithelial subpopulation with mesenchymal features aligns with a mesenchyme-like subtype of high-grade serous ovarian cancer.
- Rare basal ‘peg’ cells exhibit fetal mesonephric developmental transcriptional programs and are maintained *ex-vivo* in fallopian tube organoids.

## Introduction

High-grade serous ovarian carcinoma (HGSOC) is the most common and lethal subtype of epithelial ovarian cancer, accounting for the majority of ovarian cancer-related deaths worldwide.^1,2^ Once thought to originate from the ovarian surface epithelium, compelling molecular, genomic, and pathological evidence now supports the distal fallopian tube (FT) epithelium, particularly secretory cells of the fimbria, as the principal site of origin in most HGSOC cases.^3–9^ This paradigm shift has focused research on the pathogenesis of these cancers to the FT epithelium in an effort to develop novel early detection or interception approaches.

The FT epithelium comprises two major lineages: multiciliated cells, which direct transport of gametes and embryos through coordinated ciliary motion, and secretory cells, which are a source of cytokines, growth factors, and mucins that shape the luminal microenvironment throughout the menstrual cycle.^10–17^ Both cell lineages are hormonally regulated and exhibit region-specific specialization along the tubal axis.^11,18–25^ Increasing evidence implicates secretory cells as the predominant cell-of-origin in serous carcinogenesis, supported by their involvement in early putative precursor lesions [e.g., p53 signatures and serous tubal intraepithelial carcinomas (STICs)] and their experimental capacity for transformation.^5,6,26–33^ Emerging single-cell atlases of the FT have begun to reveal transcriptional heterogeneity within the FT epithelium, identifying discrete subpopulations within the canonical secretory and multiciliated dichotomy.^18,19,34–36^ However, the functional roles, developmental origins, and disease relevance of these subtypes remain incompletely characterized. In particular, the identity, transcriptional regulation, and lineage potential of progenitor or stem-like epithelial cells remain elusive.

To address these challenges, we established a living biobank of FT cells and organoids as a resource to provide a robust experimental infrastructure for mechanistic studies of epithelial lineage specification, progenitor dynamics, and transformation susceptibility. Organoid technologies offer a powerful platform for modeling the FT epithelium in a physiologically relevant, patient-specific context. Organoids derived from primary FT tissue recapitulate key features of epithelial organization, apicobasal polarity, and lineage composition, while enabling controlled interrogation of cell fate decisions and disease-related perturbations across diverse clinical and genetic backgrounds.^37,38^

Leveraging this organoid resource, we performed an integrative, multi-omic analysis to define the cellular architecture and regulatory landscape of the FT epithelium. By combining single-cell transcriptomics, single-nucleus multiome profiling, proteomics, secretome analysis, ultrastructural imaging, and functional assays, we generated the most comprehensive atlas of FT epithelial cell states to date. This effort resolves both canonical and previously unrecognized epithelial subtypes, defines robust multiciliated and secretory lineage signatures, and maps transcription factor networks that govern lineage identity. Among these, we identified a rare, peg-like epithelial population with dual secretory and mesenchymal features, present in our organoid system. This population shares transcriptional similarities with fetal progenitors and the mesenchymal molecular subtype of HGSOC, supporting its potential role as a cell-of-origin for this disease. Together, our findings provide a foundational resource and framework for dissecting FT epithelial biology, differentiation trajectories, and lineage-specific transformation risk.

## Results

### Establishment of a Comprehensive Living Fallopian Tube Organoid Biobank

We established a clinically and genetically annotated FT organoid biobanking pipeline to process, analyze, and cryopreserve patient-derived FT cells and organoids. Surgical resections of FT tissue and Tao brushings were obtained from consenting patients (≥18 years of age) undergoing risk-reducing salpingectomy or other surgeries including salpingectomy at Mayo Clinic Rochester. In parallel, FT tissues were also collected during autopsies at the same institution (**Figure 1A**). The FT biobank is an ongoing, prospective protocol enabling collection of annotated specimens. Here, we describe 970 FT specimens from 202 donors collected between 2018 and 2025, encompassing a diverse range of ages, demographies, medical histories, high-risk mutation statuses, exposures, and mortality outcomes (**Table 1**). The biobank primarily consists of samples from living patients (∼84%), with autopsy samples comprising ∼16%. Donor ages span from 18 to 99 years. Although the biobank covers a diverse range of samples, it predominantly represents white individuals (93.5%) which directly reflects the demographic distribution of the geographical region sampled. The most prevalent prior cancers among donors include breast, gynecologic, and skin cancers, while fibroids, endometriosis, and pelvic organ prolapse are the most common benign conditions. Only 4 patients have a prior history of ovarian cancer, and 28 patients were exposed to chemotherapy for their prior cancers. Approximately 57% of patients have documented use of personal contraception, with the majority using oral contraceptive pills.

**Figure 1.**
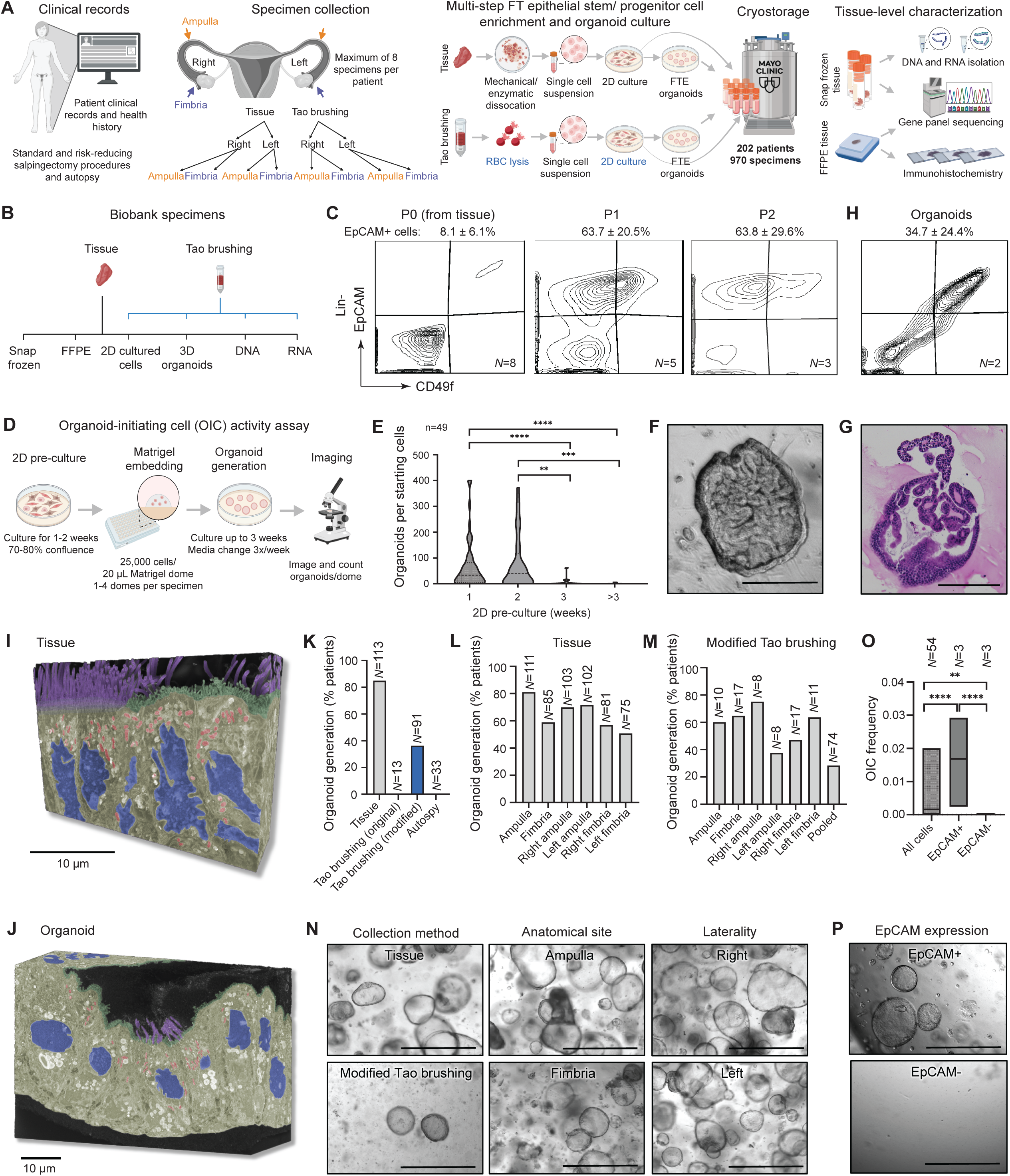
Establishment of a living fallopian tube organoid biobank reveals lineage and tissue architecture fidelity and distinct progenitor sensitivities. **(A)** Schematic overview of the biobanking workflow. **(B)** Types of biospecimens available in the biobank. **(C)** Flow cytometry analysis of EpCAM (epithelial marker) and CD49f (progenitor marker) in lineage-negative (Lin⁻: DAPI⁻, CD31⁻, CD45⁻) fallopian tube cells across multiple 2D passages (P0-P2). **(D)** Schematic of the organoid-initiating cell (OIC) activity assay workflow. **(E)** Organoid formation following different durations of 2D pre-culture (in weeks). **(F)** Representative brightfield image of a highly differentiated organoid with extensive epithelial folding and invaginations. Scale bar, 200 µm. **(G)** H&E-stained image of a representative organoid showing epithelial architecture. Scale bar, 200 µm. **(H)** Flow cytometry of EpCAM and CD49f in Lin⁻ cells from three-week-old organoids. **(I and J)** Serial block-face scanning electron microscopy (SBFSEM) 3D reconstructions of fallopian tube tissue (**I**) and a corresponding organoid (**J**). Cilia are shown in purple, microvilli in green, nuclei in blue, mitochondria in red, and cytoplasm and other organelles in yellow. **(K)** Organoid generation success rates across patients stratified by biospecimen collection method. **(L and M)** Organoid generation success rates from tissue (**L**) and modified Tao brushing (**M**) samples, further stratified by anatomical site (ampulla vs. fimbria) and laterality (right vs. left). **(N)** Representative images of organoids by collection method, anatomical site, and laterality. Scale bars, 1000 µm. **(O)** OIC frequencies of unseparated passage 1 (P1) cells and FACS-sorted EpCAM⁺ and EpCAM⁻ Lin^-^ subsets. **(P)** Representative images of organoids derived from EpCAM⁺ and EpCAM⁻ cells. Scale bars, 1000 µm. ***N***=patients; **n**=specimens (applies to all panels). Significance was determined by Mann-Whitney test; **, *p* < 0.01; ***, *p* < 0.001; ****, *p* < 0.0001. ***See Supplementary*** Figures 1 and 2 ***for additional related data*.**

**Table 1.** Patient characteristics. * indicates more than one category is possible per patient. The ‘Other’ category for race reflects participants who did not identify with the predefined race options in the Mayo Clinic medical record and instead selected ‘Other’. Other cancers include musculoskeletal, respiratory/thoracic, genitourinary, and endocrine. Other reproductive diseases include ovarian borderline tumor, cervical dysplasia, benign ovarian neoplasm, and malignant ovarian neoplasm. Other reproductive surgeries include ovarian cystectomy, unilateral salpingectomy, labioplasty, endometrial biopsy, endometrial ablation, loop electrosurgical excision procedure (LEEP), cystoscopy, unilateral oophorectomy, and endometrial excision. Other chronic diseases include but are not limited to chronic kidney disease, autoimmune disorders and more. COVID-19 vaccination and SARS-CoV-2 exposure data were not assessed for specimens collected before 2020, as these events predated vaccine availability and widespread viral circulation. Abbreviations: RRSO, risk-reducing salpingo-oophorectomy; PCOS; polycystic ovarian syndrome; C section; caesarean section; IUD, intrauterine device; IVF *in-vitro* fertilization.

| Characteristics | Total patients (N=202) | Patients with mutation (N=33) |
| --- | --- | --- |
|  | No. (%) | No. (%) |
| **Vital status at tissue collection** |  |  |
| Living | 169 (83.66) | 32 (96.97) |
| Deceased | 33 (16.34) | 1 (3.03) |
| **Age (years)** |  |  |
| <45 | 75 (37.13) | 17 (51.52) |
| 45-60 | 72 (35.64) | 10 (30.30) |
| >60 | 55 (27.23) | 6 (18.18) |
| **Race** |  |  |
| White | 189 (93.55) | 32 (96.97) |
| Black/ African American | 1 (0.50) | 0 (0.00) |
| Native American/ American Indian | 1 (0.50) | 0 (0.00) |
| Asian/ Pacific Islander | 5 (2.48) | 0 (0.00) |
| Other | 2 (0.99) | 1 (3.03) |
| Data missing | 4 (1.98) | 0 (0.00) |
| **Surgical method** |  |  |
| Salpingectomy | 34 (16.83) | 2 (6.06) |
| RRSO | 21 (10.40) | 15 (45.46) |
| Hysterectomy + RRSO | 10 (4.95) | 6 (18.18) |
| Hysterectomy + salpingectomy | 73 (36.14) | 2 (6.06) |
| Hysterectomy + salpingo-oophorectomy | 16 (7.92) | 3 (9.09) |
| Salpingo-oophorectomy | 15 (7.42) | 4 (12.12) |
| Autopsy | 33 (16.34) | 1 (3.03) |
| **Gravidity** |  |  |
| 0 | 29 (14.36) | 4 (12.12) |
| 1 | 15 (7.42) | 3 (9.09) |
| 2 | 51 (25.25) | 8 (24.24) |
| 3 | 36 (17.82) | 8 (24.24) |
| >3 | 37 (18.32) | 9 (27.28) |
| Data missing | 34 (16.83) | 1 (3.03) |
| **Parity** |  |  |
| 0 | 36 (17.82) | 5 (15.15) |
| 1 | 22 (10.89) | 4 (12.12) |
| 2 | 61 (30.20) | 7 (21.22) |
| 3 | 29 (14.36) | 8 (24.24) |
| >3 | 26 (12.87) | 8 (24.24) |
| Data missing | 28 (13.86) | 1 (3.03) |
| **Menopausal status** |  |  |
| Premenopausal | 76 (37.62) | 14 (42.43) |
| Perimenopausal | 8 (3.96) | 2 (6.06) |
| Postmenopausal | 75 (37.13) | 12 (36.36) |
| Data missing | 43 (21.29) | 5 (15.15) |
| **Age at menarche (years)** |  |  |
| <10 | 2 (0.99) | 0 (0.00) |
| 10-16 | 90 (44.55) | 25 (75.76) |
| >16 | 4 (1.98) | 1 (3.03) |
| Data missing | 106 (52.48) | 7 (21.21) |
| **Mutation carrier** |  |  |
| Yes | 33 (16.34) | 33 (100.00) |
| No | 31 (15.35) | 0 (0.00) |
| Data missing | 138 (68.31) | 0 (0.00) |
| **History of cancer\*** |  |  |
| Breast | 27 (13.37) | 12 (36.36) |
| Gynecologic | 11 (5.45) | 0 (0.00) |
| Skin | 10 (4.95) | 3 (9.09) |
| Hematologic/ blood | 5 (2.48) | 1 (3.03) |
| Digestive/ gastrointestinal | 3 (1.49) | 2 (6.06) |
| Other | 4 (1.98) | 1 (3.03) |
| Data missing | 149 (73.76) | 17 (51.52) |
| **History of ovarian cancer** |  |  |
| Yes | 4 (1.98) | 0 (0.00) |
| Data missing | 198 (98.02) | 33 (100.00) |
| **History of chemotherapy** |  |  |
| Yes | 30 (14.85) | 14 (42.42) |
| Data missing | 172 (85.15) | 19 (57.58) |
| **History of reproductive disease\*** |  |  |
| Fibroids | 64 (31.68) | 13 (39.39) |
| Endometriosis | 35 (17.33) | 2 (6.06) |
| Pelvic organ prolapse | 27 (13.37) | 23 (69.70) |
| Endometrial hyperplasia | 7 (3.47) | 0 (0.00) |
| PCOS | 7 (3.47) | 1 (3.03) |
| Other | 4 (1.98) | 0 (0.00) |
| Data missing | 82 (40.59) | 17 (51.52) |
| **History of reproductive surgeries\*** |  |  |
| Dilation and curettage | 35 (17.33) | 5 (15.15) |
| C Section | 29 (14.36) | 6 (18.18) |
| Hysterectomy | 17 (8.42) | 6 (18.18) |
| Tubal ligation | 16 (7.92) | 2 (6.06) |
| Myomectomy | 10 (4.95) | 3 (9.09) |
| Elective abortion | 9 (4.46) | 1 (3.03) |
| Hysteroscopy | 9 (4.46) | 1 (3.03) |
| Polypectomy | 5 (2.48) | 0 (0.00) |
| Other | 24 (11.88) | 1 (3.03) |
| Data missing | 92 (45.54) | 15 (45.45) |
| **Personal contraception** |  |  |
| Yes | 121 (59.90) | 24 (72.73) |
| No | 4 (1.98) | 0 (0.0) |
| Data missing | 77 (38.12) | 9 (27.27) |
| **Type of personal contraception** |  |  |
| Pill | 49 (24.24) | 11 (33.33) |
| Hormonal IUD | 11 (5.45) | 1 (3.03) |
| IUD | 9 (4.46) | 3 (9.09) |
| Tubal ligation | 5 (2.48) | 0 (0.00) |
| Injection | 3 (1.49) | 1 (3.03) |
| Implant | 1 (0.50) | 1 (3.03) |
| Patch | 1 (0.50) | 1 (3.03) |
| Diaphragm | 1 (0.50) | 0 (0.00) |
| Multiple methods | 39 (19.31) | 5 (15.15) |
| Data missing | 83 (41.07) | 10 (30.31) |
| **Infertility treatment** |  |  |
| IVF | 11 (5.45) | 1 (3.03) |
| Repeated ovulation injections | 1 (0.5) | 1 (3.03) |
| None | 19 (9.41) | 2 (6.06) |
| Data missing | 171 (84.64) | 29 (87.88) |
| **Chronic disease\*** |  |  |
| Hypertension | 41 (20.30) | 7 (21.21) |
| Hyperlipidemia | 25 (12.38) | 2 (6.06) |
| Diabetes | 11 (5.45) | 1 (3.03) |
| Other | 65 (32.18) | 14 (42.42) |
| Data missing | 65 (32.18) | 12 (36.36) |
| **Body-mass index** |  |  |
| <25 | 52 (25.74) | 10 (30.30) |
| 25-29 | 52 (25.74) | 5 (15.15) |
| 30-39 | 57 (28.22) | 13 (39.40) |
| ≥40 | 32 (15.84) | 4 (12.12) |
| Data missing | 9 (4.46) | 1 (3.03) |
| **History of HPV infection** |  |  |
| Yes | 22 (10.89) | 1 (3.03) |
| No | 124 (61.39) | 19 (57.58) |
| Data missing | 56 (27.72) | 13 (39.39) |
| **Smoking** |  |  |
| Active | 10 (4.95) | 1 (3.03) |
| History | 49 (24.25) | 9 (27.27) |
| Never | 142 (70.30) | 23 (69.70) |
| Data missing | 1 (0.5) | 0 (0.00) |
| **Alcohol consumption** |  |  |
| Active | 133 (65.84) | 27 (81.82) |
| History | 27 (13.37) | 3 (9.09) |
| Never | 38 (18.81) | 2 (6.06) |
| Data missing | 4 (1.98) | 1 (3.03) |
| **COVID exposure** |  |  |
| Yes | 40 (19.80) | 8 (24.24) |
| No | 69 (34.16) | 15 (45.46) |
| Data missing | 93 (46.04) | 10 (30.30) |
| **COVID vaccine** |  |  |
| Yes | 88 (43.56) | 15 (45.46) |
| No | 15 (7.43) | 7 (21.21) |
| Data missing | 99 (49.01) | 11 (33.33) |

Patients with a family history of genetic mutations, breast or ovarian cancer, and reproductive diseases are often offered genetic sequencing to identify germline mutations associated with high cancer risk, for which corresponding clinical genetic testing records are available for biobank patients **(Supplementary Figure S1A)**.^39^ From the electronic medical record (EMR), 33 patients were identified with high-risk mutations (∼16% of the total patients) **(Supplementary Figure S1B)**. The most common mutations were those associated with Hereditary Breast and Ovarian Cancer syndrome (HBOC), specifically *BRCA1* and *BRCA2* **(Supplementary Figure S1C)**. Other prominent mutations include those in DNA mismatch repair genes, *MSH2*, *MSH6*, and *MLH1*, which are linked to Lynch syndrome. Notably, four patients were found to carry double mutations. Hierarchical clustering was used to explore associations between mutations, vital status, race, age, menopausal status, and cancer history; however, no strong associations were observed in this cohort **(Supplementary Figure S1D)**.

For each patient, a maximum of 8 specimens were collected and tracked using Mayo Clinic’s Research Laboratory Information Management System (**Figure 1A**). We processed samples using a protocol,^40^ previously modified from Kessler et al,^37^ that includes collection of resected FT tissue from the Frozen Section Laboratory, enzymatic dissociation, manual mechanical disaggregation with a scalpel, epithelial-rich cell isolation, and 2D pre-culture prior to 3D organoid generation in a Matrigel culture system (**Figure 1A**). Tao brushings were obtained intraoperatively following FT removal. Specifically, a brush was repeatedly inserted into the resected FTs to collect cells with direct interface and proximity to the lumen **(Supplementary Figure S2A)**. The brushes were submerged in collection media and transported to the research laboratory. Subsequently, the cells were collected by agitation of brushes in the media and centrifugation. Red blood cell lysis was performed prior to culture in 2D and subsequently in 3D for organoid generation and cryopreservation (**Figure 1A**). In addition to cryopreservation of all cultured cells, samples were processed for nucleic acid extraction (DNA and RNA) and snap frozen for future studies. Also, access to clinical formalin-fixed paraffin-embedded (FFPE) tissue was provided by the Mayo Clinic tissue registry (**Figure 1A-B**).

### Tubal Organoids Recapitulate Lineage Differentiation and Tissue Epithelial Architecture

We evaluated freshly isolated single cells [passage 0 (P0)] or 2D culture expanded (P1 and P2) cells for their epithelial content via flow cytometry using the epithelial cell surface marker, EpCAM, and progenitor marker, CD49f.^41^ EpCAM and CD49f have been extensively validated for separation of epithelial populations in other tissues including mammary gland^42^, prostate^43^, tonsil^44^, and salivary gland^45^. Our optimized 2D culture conditions progressively enriched EpCAM+ epithelial cells with successive passages (**Figure 1C**). These epithelial-enriched cells were then used to generate organoids via an organoid-initiating cell (OIC) activity assay. Patient cells were pre-cultured in 2D for 1-2 weeks until they reached 70-80% confluence, after which 25,000 P1 cells were embedded in 20 uL Matrigel domes (**Figure 1D**). The duration of 2D pre-culture was restricted to 1-2 weeks because cells maintained in pre-culture for 3 or more weeks showed a marked reduction in organoid formation, suggesting loss of progenitor function (**Figure 1E**). Additional culture and processing conditions, including dissociation duration, treatment of culture plates with collagen, and incubator oxygen content, were optimized to maximize organoid formation **(Supplementary Figure S2B-D)**. Further, cryopreservation of cells and organoids did not diminish organoid forming capacity **(Supplementary Figure S2E)**. With the optimized culture conditions, we reliably generated FT organoids that closely resembled those described in previous reports.^20,37,38^ These organoids maintained a spherical, lumenized structure with epithelial folding, mirroring the architecture of native FT tissue (**Figure 1F-G**).^37^ Additionally, organoids retained a high percentage of EpCAM+ cells following 3 weeks in culture (**Figure 1H**).

To gain ultrastructural insight into organoid morphology and cell composition, we performed serial block-face scanning electron microscopy (SBFSEM) on FT tissue and organoids. This high-resolution 3D imaging approach enabled detailed visualization of epithelial architecture, confirming the presence of both multiciliated and secretory cells within the organoids (**Figure 1I-J, Supplementary Figure S2F)**. Multiciliated cells were identified by dense tufts of apical motile cilia projecting into the lumen, while secretory cells displayed characteristic luminal microvilli and apical secretory vesicles, closely resembling their *in-vivo* counterparts **(Supplementary Figure S2G)**. The spatial arrangement and organization of these cell types mirrored those observed in native FT epithelium, including epithelial polarity and lumen-facing orientation **(Supplementary Figure S2H)**. These findings support the structural fidelity of the organoid model and reinforce its utility for studying FT epithelial biology in a physiologically relevant context.

### Characterization of Factors Influencing Tubal Organoid Generation Identifies Autopsy Tissue as Unfit for Growth, Revealing Unique Progenitor Sensitivity

To evaluate whether the method of collection influenced organoid formation, we quantified organoid generation by patient. Tissue samples yielded the highest success rate, with organoids generated in ∼83% of cases (**Figure 1K**). At the time of biobank initiation, Tao brushing was an exploratory method requiring procedural refinement. Early samples were obtained intraoperatively, transported in collection media, and embedded directly in Matrigel for culture without further processing (**Figure 1A**). However, this method failed to generate organoids in the first 13 Tao brushing collections **(Supplementary Figure S2I)**. Protocol refinement, including red blood cell lysis and a 2D pre-culture step, improved Tao brushing outcomes, enabling organoid generation in ∼37% of patients (**Figure 1K**). In stark contrast, single cells isolated from autopsy-derived FT tissue (*N*=33) uniformly failed to expand or form organoids, reflecting severely compromised progenitor fitness incompatible with *ex-vivo* culture. This finding diverges from reports in other epithelial systems, such as mammary gland, where autopsy tissue reliably supports organoid generation, underscoring a unique vulnerability of FT progenitors to postmortem degradation.^45^

Next, we evaluated organoid generation stratified by anatomical site and laterality in tissue specimens. We found that samples of the ampulla yielded a higher frequency of organoid formation compared to the fimbria, independent of laterality (**Figure 1L**). In Tao brushing specimens, organoid generation rates were similar between the ampulla and fimbria but displayed greater variability when analyzed by laterality (**Figure 1M**). Upon receiving tissue samples, specimen weights were recorded, allowing us to assess organoid generation relative to tissue weight. Among specimens that successfully formed organoids, organoid yield per milligram of tissue followed a normal distribution **(Supplementary Figure S2J)**. Ampulla specimens produced slightly more organoids per milligram of tissue than fimbria, while no significant difference was observed between left and right specimens or specimens from mutation carriers and non-carriers **(Supplementary Figure S2K-M)**. Altogether we show that organoids can be generated independent of anatomical site, laterality, collection method, or mutation status, patient-specific heterogeneity is observed in organoid formation across different sites, lateralities, and collection methods (**Figure 1N; Supplementary Figure S2N-P)**.

Overall, the frequency of organoid initiating cells in our system ranges from approximately <0.000004 to 0.029, with a mean of 0.0016 (**Figure 1O**). Flow cytometry previously demonstrated that EpCAM+ cells are enriched through successive passages in 2D culture (**Figure 1C**), and we show that these pure, sorted EpCAM+ cells are responsible for 100% of the organoid-initiating activity, as EpCAM-cells fail to generate organoids (**Figure 1O-P**). In some patient samples, EpCAM⁻ cells give rise to fibrotic outgrowths in Matrigel, likely originating from stromal cells within the non-epithelial compartment **(Supplementary Figure S2Q)**.

### Multi-omics Approach Establishes Robust Tubal Epithelial Lineage-specific Signatures

To comprehensively define the cellular and molecular landscape of the FT epithelium, we employed a systematic multi-omics framework leveraging both freshly dissociated tissues and biobank-derived organoids. We performed single-cell RNA sequencing (scRNA-Seq), single-nucleus RNA and ATAC sequencing (snMultiome), and bulk RNA-Seq on freshly dissociated FT cells and pooled organoids (**Figure 2A**). To broaden the cellular and patient diversity of our analysis, we integrated both newly generated and publicly available FT single-cell atlases, representing both normal and high-risk mutant samples (**Figure 2B**). Our in-house datasets included individuals with *BRCA2* and *TP53/BRCA2* germline mutations, while public data was used to expand the cohort to include *BRCA1*-mutant samples.^18,19,34,35,46^ Dataset integration was performed using Harmony-based batch correction to align shared cell states across patients, batches, and mutation backgrounds while preserving biological variance. Together with our in-house datasets, this enabled the construction of the largest known integrated FT atlas encompassing 256,779 cells and 19,419 protein-coding genes from 43 patients.

**Figure 2.**
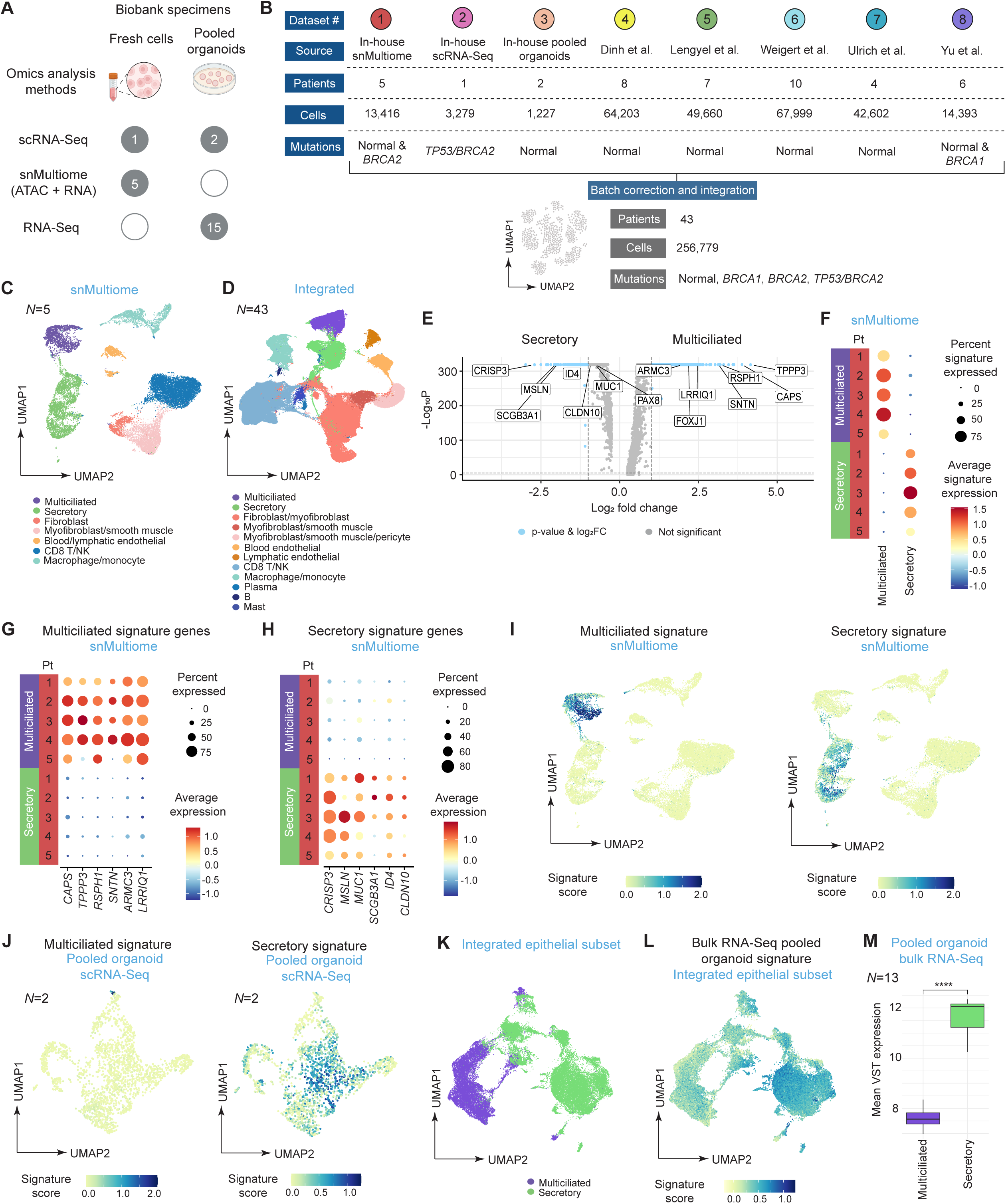
Multi-omics Approach Establishes Robust Tubal Epithelial Lineage-specific Signatures. **(A)** Summary of nucleic acid-based omics technologies applied to biobank specimens. The number of specimens analyzed per modality are indicated. **(B)** Schematic overview of in-house and public single-cell RNA sequencing (scRNA-seq) and single-nucleus multiome (snMultiome) datasets used to generate the integrated FT atlas. **(C)** UMAP showing cell type annotations from the snMultiome datasets. **(D)** UMAP showing cell type annotations in the integrated dataset. **(E)** Volcano plot of multiciliated vs. secretory lineages in integrated dataset. **(F)** Application of the multiciliated and secretory lineage signatures to lineage-specific cells from the snMultiome dataset, shown by patient (*N*=5). **(G and H)** Expression of the individual genes comprising the multiciliated **(G)** and secretory **(H)** signatures in multiciliated and secretory cells from the snMultiome dataset, split by patient. **(I)** Application of multiciliated and secretory lineage signatures to the snMultiome UMAP, showing distribution across all cell types. **(J)** UMAP of pooled organoid samples showing application of the multiciliated and secretory lineage signatures. **(K)** UMAP of the epithelial subset from the integrated dataset, labeled by epithelial lineage. **(L)** UMAP of the integrated epithelial subset showing the pooled organoid RNA-seq signature, based on the top 200 epithelial genes shared by ≥10 samples. **(M)** Average expression of both lineage signatures across 15 pooled organoid bulk RNA-seq datasets (*N*=13; n=15). ***N*=**patients; **n**=specimens (applies to all panels). Significance was determined by two-sample t-test; ****, *p* < 0.0001. ***See Supplementary*** Figure 3 ***for additional related data*.**

Clustering analysis of the snMultiome data individually identified 7 distinct clusters, corresponding to known FT cell types including multiciliated and secretory lineage epithelial cells, fibroblasts, myofibroblasts, smooth muscle cells, blood and lymphatic endothelial cells, CD8 T cells, NK cells, macrophages and monocytes (**Figure 2C**). Integration with additional scRNA-seq datasets further enhanced resolution and revealed additional cell populations not consistently captured by individual datasets, including mast cells, plasma cells, B cells, and pericytes (**Figure 2D**).

Although prior studies have characterized epithelial lineages in the FT, reported marker genes for multiciliated and secretory cells have shown limited consistency in expression and abundance across datasets and platforms, likely reflecting differences in tissue processing, patient cohorts, and analytic pipelines.^18,19,34,35^ In addition, substantial intra-lineage heterogeneity in currently defined lineage marker expression has been observed, complicating the use of these markers in functional studies aimed at isolating or characterizing specific epithelial subsets. Specifically, the multiciliated lineage marker, *FOXJ1*, and the secretory lineage marker, *PAX8*, show variable abundance, lineage specificity and expression across datasets **(Supplementary Figure S3A)**. To address these limitations and enable reproducible identification of FT epithelial subtypes across modalities, we sought to define robust lineage-specific gene signatures applicable to both scRNA-seq and snMultiome datasets. First, canonical markers were used to guide the initial annotation of multiciliated and secretory epithelial clusters. We then performed differential gene expression analysis between these lineages across individual in-house and public datasets (**Figure 2E**). Genes that were consistently enriched in one lineage across multiple datasets were retained and systematically assessed for their expression patterns across all cells within both lineages. To ensure robustness, we prioritized genes with uniform and prominent intra-lineage expression and high lineage specificity. These refined gene sets were compiled into lineage-specific signatures and validated in independent public FT datasets and the integrated dataset **(Supplementary Figure S3B-C)**.

This analysis defined two robust 6-gene signatures marking multiciliated (*CAPS, TPPP3, RSPH1, SNTN, ARMC3, LRRIQ1*) and secretory (*CRISP3, MSLN, MUC1, SCGB3A1, ID4, CLDN10*) lineages, which, when applied as composite markers, reliably and specifically distinguished each lineage from all other cell types across patients **(Supplementary** Figure 3D-E). These signatures represent novel marker combinations that include both previously characterized lineage genes and genes previously unannotated in the context of FT epithelial identity. The signatures and genes within each signature showed high lineage specificity, with consistent expression across individual patients in the snMultiome dataset (**Figure 2F-I**). Gene signatures, in general, provide a more reliable measure of lineage identity by combining the expression of multiple co-expressed markers, improving signal strength and reducing single-gene variability. By leveraging multi-gene expression, they offer a biologically robust and reproducible framework for distinguishing cell lineages, while minimizing inter-patient variability and enhancing specificity. Protein-level evaluation using the Human Protein Atlas confirmed detectable expression in a lineage-specific manner for 8 of the 12 signature genes in the FT epithelium, while the remaining were unreported, likely due to limited antibody availability or lack of assessment rather than true absence **(Supplementary Figure S3F-G)**.

### Multi-level Transcriptomic Analysis of Tubal Organoids Identifies Predominant Secretory Lineage Differentiation

To assess epithelial lineage composition in our organoid model, we applied the multiciliated and secretory signatures to scRNA-seq data from pooled organoids. The analysis revealed marked enrichment of the secretory signature across the majority of cells, indicating a predominance of secretory cells in the organoids, consistent with SBFSEM findings (**Figure 2J; Supplementary** Figure 3H; **Figure 1I-J**). To explore this further, we derived an epithelial organoid signature from the top 200 epithelial genes shared by ≥10 samples and projected it onto the integrated epithelial subset, where it was predominantly expressed in secretory clusters, reinforcing the secretory identity of the organoids (**Figure 2K-L; Supplementary Figure S3I)**. Finally, we applied lineage signatures to bulk RNA-seq data from 15 pooled organoid specimens (*N*=13; n=15), combining anatomical sites and lateralities based on the absence of significant transcriptomic differences across these variables **(Supplementary Figure S3J)**. Application of both lineage signatures in this setting further confirmed heightened expression of the secretory lineage signature in organoids, consistent with scRNA-seq results and supporting the robustness of these signatures across fresh tissue and organoid contexts (**Figure 2M**).

### Transcriptional Regulation and Gene Regulatory Networks Define Lineage Identity and Heterogeneity in Tubal Epithelium

Distinct transcription factor (TF) programs define epithelial lineage identity in the FT. ChromVAR^47^ analysis of snMultiome data revealed enrichment of *RFX* and *FOX* TF families in multiciliated cells, while *SOX* and *TEAD* families dominated secretory cells (**Figure 3A**). Shared TFs including *NFIX, NFIC, SNAI1, HNF1B,* and *ZEB1* showed activity across both lineages, although they exhibited differing patterns of target gene regulation **(Supplementary Figure S4A-B).**

**Figure 3.**
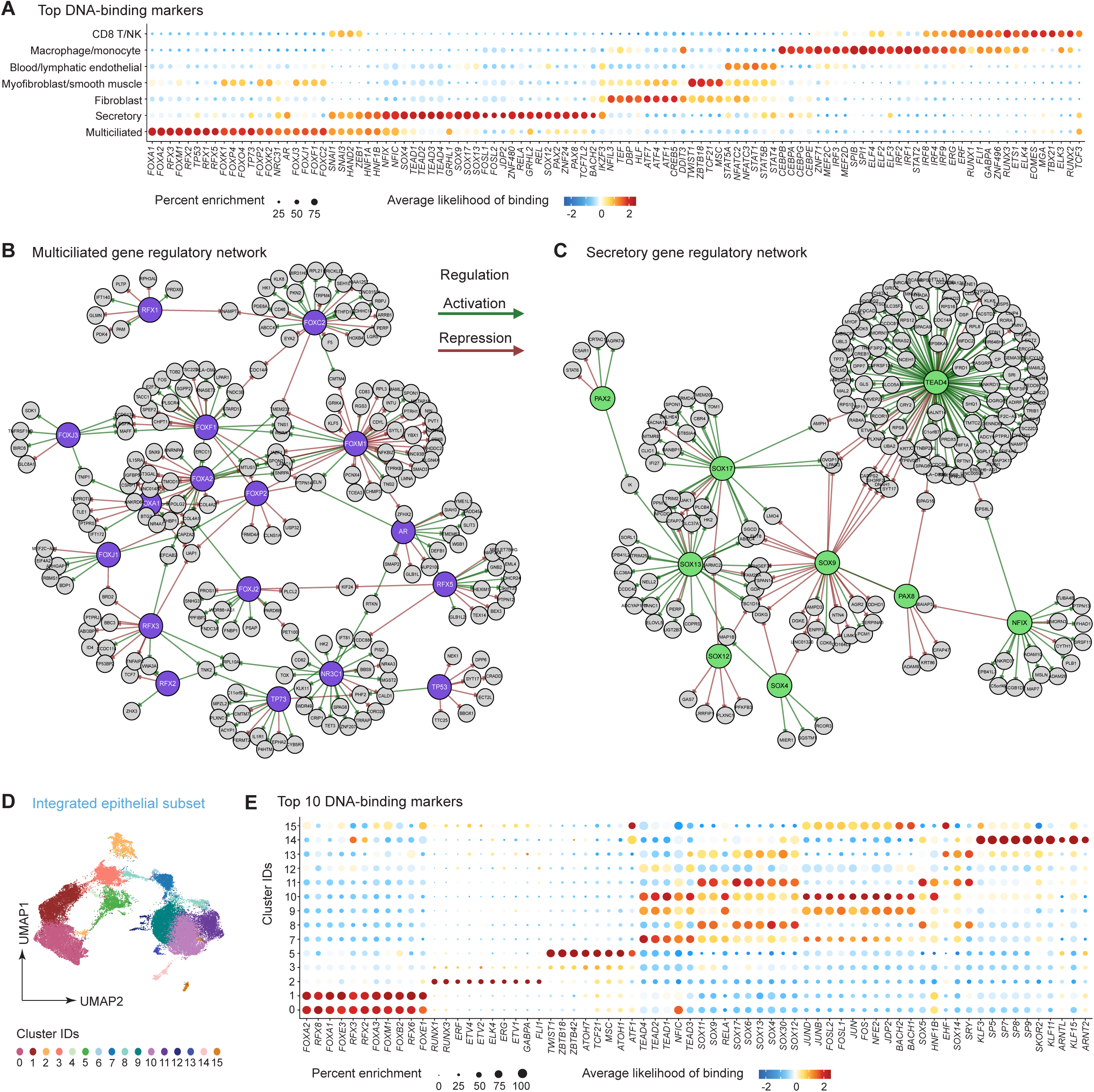
Transcriptional Regulation and Gene Regulatory Networks Define Lineage Identity and Heterogeneity in Tubal Epithelium. **(A)** Top DNA binding factor markers for each cell type predicted by ChromVAR analysis (DNA binding factor score) in snMultiome dataset. **(B and C)** Gene regulatory networks (GRNs) inferred by FigR for multiciliated **(B)** and secretory **(C)** cells in the snMultiome dataset. Green arrows indicate activation of target genes; red arrows indicate repression. Purple and green central nodes represent transcription factors (TFs), while gray peripheral nodes represent downstream TF targets. **(D)** UMAP of the integrated epithelial subset, colored by transcriptionally defined subclusters (0–15). **(E)** Top 10 DNA binding factor markers per epithelial subcluster predicted by ChromVAR analysis from the snMultiome dataset. ***See Supplementary*** Figure 4 ***for additional related data*.**

Using FigR^48^, we constructed lineage-specific gene regulatory networks, uncovering that *RFX3* and related TFs drive canonical ciliogenesis genes (e.g. *CCDC114*, *EFCAB2*) in multiciliated cells while repressing the secretory marker *ID4* (**Figure 3B**). In secretory cells, *SOX17* acts as a key activator promoting *OVGP1*, whereas SOX9 primarily functions as a repressor. Notably, *SOX9* interacts with *TEAD3*, *RELA*, and *REL* to activate *PAX8*, indicating a broader, TF-driven regulatory program governing this critical secretory lineage marker (**Figure 3C**). Additional regulators such as *NFIX* and *TEAD4* showed both activating and repressive functions: *NFIX* activated *MSLN*, a secretory lineage signature gene, while *TEAD4* activated *CLDN4*, a key epithelial structural gene, and repressed *OVGP1* in coordination with *SOX9*. Further, multiple TFs—*BACH2*, *FOSL1*, and *RELA*— activate *MUC1*, a transmembrane glycoprotein with roles in establishing a FT protective mucosal barrier and also a component of the secretory lineage signature (data not shown). In addition, a combination of TFs activate *MUC16* (CA125), a membrane-associated glycoprotein,^49^ and *WFDC2* (HE4), a secreted glycoprotein,^50^ which are both widely used as biomarkers for HGSOC, though limited in their effectiveness for early detection. To validate lineage-specific expression of selected transcription factors, we leveraged the Human Protein Atlas. RFX3 exhibited specificity for multiciliated cells, while SOX9 was enriched in secretory cells **(Supplementary Figure S4C)**.

To investigate intra-epithelial heterogeneity, we subclustered the epithelial compartment of the integrated dataset, revealing 16 transcriptionally distinct clusters: 2 multiciliated (clusters 0 and 1), 11 secretory (clusters 4, 6-15), and 3 mixed-lineage (clusters 2,3,5) (**Figure 3D**; **Figure 2K**). Clusters 4 and 6 were present in scRNA-Seq datasets but were not captured in the snMultiome dataset, precluding inference of their transcriptional regulators **(Supplementary Figure S4D)**. While most clusters were represented across patients, with variable relative abundance **(Supplementary Figure S4D–E)**, a subset displayed marked enrichment in FT samples from mutation carriers **(Supplementary Figure S4F)**. Specifically, *BRCA1*-mutant patients were enriched in cluster 12, *BRCA2*-mutant patients enriched in cluster 7, and the *TP53/BRCA2* double-mutant patient enriched in clusters 2, 3 and 13.^51^ This enrichment highlights possible early lineage shifts or epithelial state changes associated with germline risk, underscoring their potential relevance to predisposition or early tumorigenesis. To further contextualize these and other epithelial populations, we compared all subclusters to those reported in individual published studies.^18,34,35^ Many clusters showed strong overlap with previously defined populations **(Supplementary Figure S4G)**, reinforcing the validity of our classification across platforms and cohorts. Importantly, multiciliated clusters 0 and 1 exhibited strong association with multiple ciliated clusters, secretory clusters 9 to 15 showed great association with secretory-like clusters, and cluster 5 exhibited very strong association with subclusters of both lineages. Overall, multiciliated clusters shared a conserved TF program, whereas secretory clusters displayed greater TF diversity, suggesting increased regulatory heterogeneity within the secretory lineage (**Figure 3E**).

### Aging Alters Tubal Epithelial Composition while Preserving Progenitor Function

To investigate aging effects on FT epithelial dynamics, we compared epithelial and lineage-negative (Lin⁻) non-epithelial cells, including fibroblasts, myofibroblasts, smooth muscle cells, and pericytes, across patients spanning a wide age range using our integrated scRNA-seq dataset (**Figure 4A**). Epithelial-to-Lin⁻ cell ratios declined sharply with age, reflecting a relative loss of epithelial cells during aging (**Figure 4B**). Within the epithelium, the ratio of multiciliated to secretory cells declined significantly with age, indicating a lineage shift favoring secretory cells over time (**Figure 4C**). These findings align with previous histological studies reporting age-related reductions in multiciliated cell abundance.^52^

**Figure 4.**
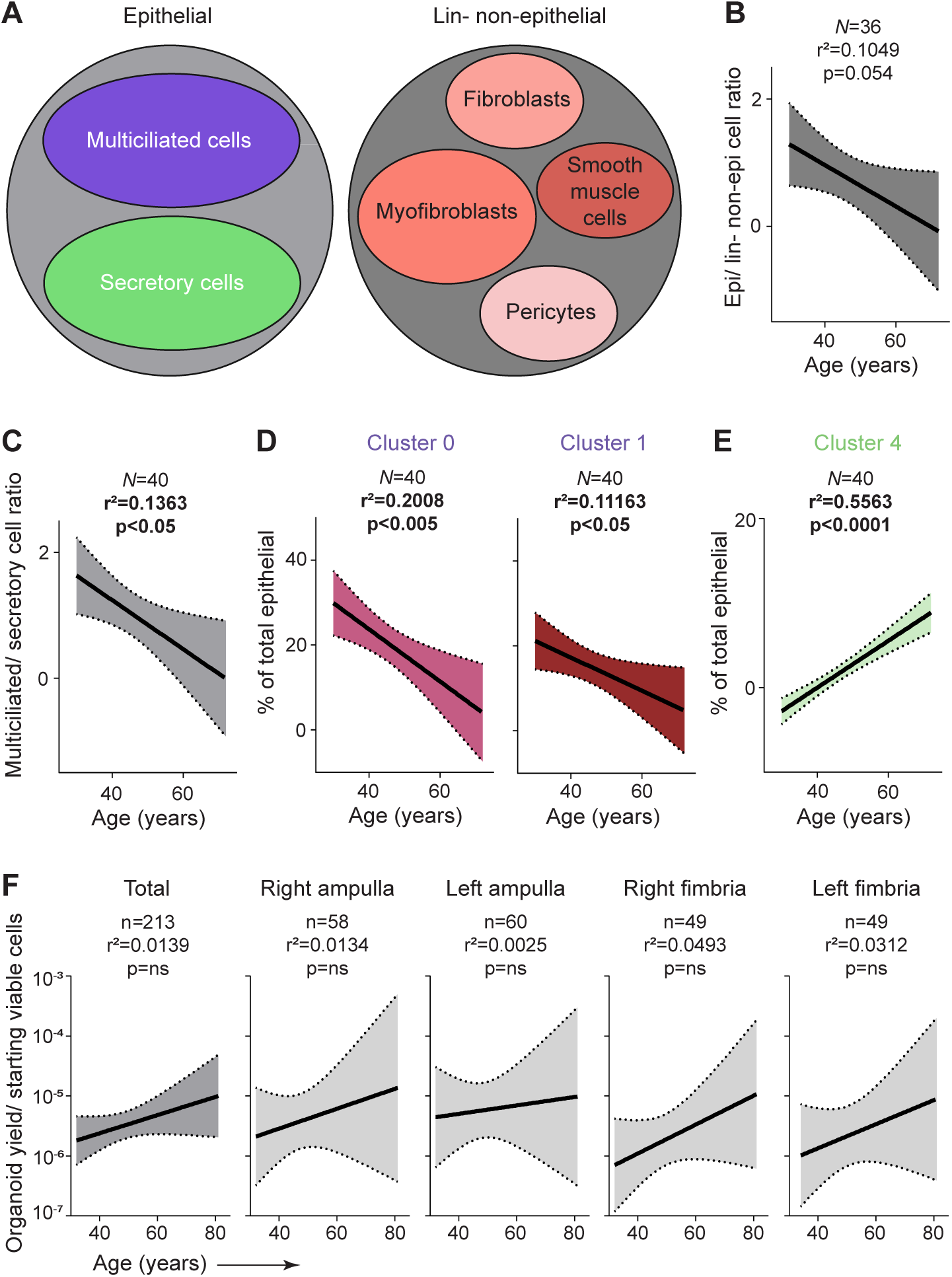
Aging Alters Tubal Epithelial Composition while Preserving Progenitor Function. **(A)** Schematic illustrating the cellular composition of the fallopian tube epithelium and the lineage-negative (Lin⁻: DAPI⁻, CD31⁻, CD45⁻) non-epithelial stromal fraction. **(B)** Correlation between the epithelial-to-Lin⁻ non-epithelial cell ratio and patient age. **(C)** Correlation between the multiciliated-to-secretory cell ratio and patient age. **(D and E)** Correlation between the contribution of specific epithelial subclusters to the total epithelial compartment and patient age: multiciliated clusters 0 and 1 (**D**), and secretory cluster 4 (**E**). **(F)** Correlation between the number of organoids generated per starting number of viable cells and patient age, shown for all patients and stratified by anatomical site (ampulla vs. fimbria) and laterality (right vs. left). ***N=*** patients; **N=** specimens (applies to all panels). Significance was determined by simple linear regression; ns, not significant. Dotted lines represent 95% confidence interval. ***See Supplementary*** Figure 5 ***for additional related data*.**

To resolve these changes in greater detail, we examined how patient age correlates with the relative abundance of transcriptionally defined epithelial subclusters within the total epithelium (**Figure 4D-E; Supplementary Figure S5A)**. Multiciliated clusters 0 and 1 declined significantly with age, with cluster 0 showing the strongest effect (**Figure 4D**). In contrast, secretory cluster 4 increased with age and was restricted to postmenopausal individuals, suggesting it represents an age-associated postmenopausal subpopulation (**Figure 4E**).

To determine whether epithelial progenitor function changed with age, we assessed organoid-initiating capacity. Despite an age-related decline in total viable epithelial cells **(Supplementary Figure S5B)**, organoid-forming efficiency was preserved (**Figure 4F**), indicating that regenerative potential remains intact with aging and menopause.

### Secretory Profiles Identify Epithelial Subpopulation–Specific Biological Function in the Fallopian Tube

The FT epithelium produces a complex array of secreted molecules that maintain epithelial integrity, barrier function, and mucosal immunity, while also modulating the luminal environment to support fertilization and embryo transport. However, the precise composition and cellular sources of these factors remain poorly defined.^53^

To define secretory molecules produced by the FT epithelium, we applied a targeted multi-omics approach across biobank-derived specimens, including patient-matched fresh tissue secretomes, early-passage (P0) 2D culture secretomes, FACS-sorted EpCAM⁺ epithelial cell proteomes, and pooled organoid proteomes. To expand our analysis, we integrated publicly available datasets, including FT lavage secretomes, PAX8⁺ 2D culture secretomes, and whole-tissue proteomes (**Figure 5A**).^54–56^

**Figure 5.**
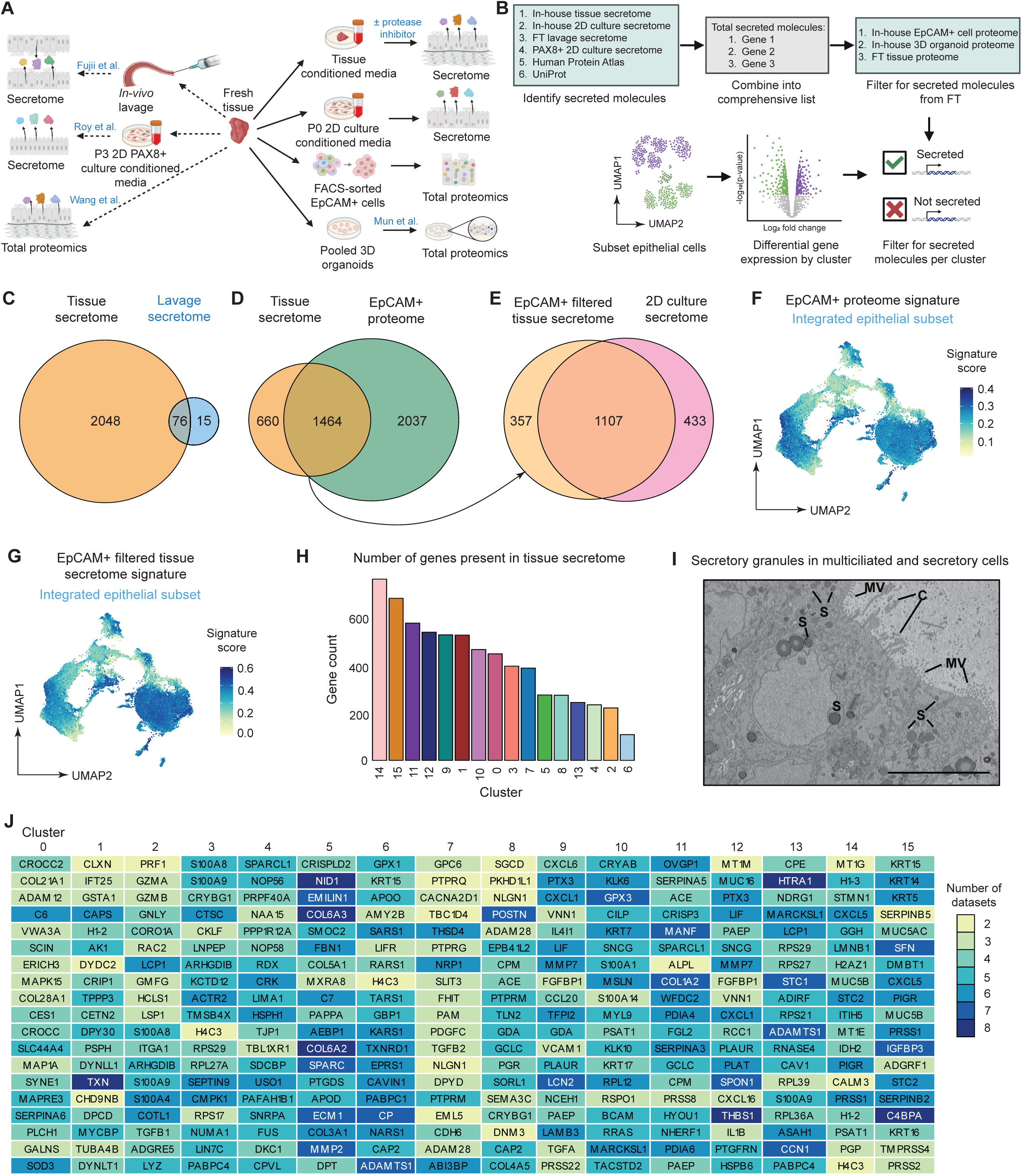
Secretory Profiles Identify Epithelial Subpopulation–Specific Biological Function in the Fallopian Tube. **(A)** Overview of proteomic and secretome datasets generated from biobank specimens and utilized from public sources. **(B)** Experimental workflow for identifying secreted molecules across the FT epithelium and their subclusters. **(C and D)** Venn diagrams comparing protein overlap between datasets: tissue secretome vs. public FT lavage secretome (**C**), and tissue secretome vs. EpCAM⁺ Lin^-^ cell proteome (**D**). **(E)** Venn diagram showing overlap between the inferred EpCAM⁺ Lin^-^ cell secretome (shared proteins from tissue secretome and EpCAM⁺ proteome) and the secretome of 2D-cultured cells. **(F)** UMAP of the integrated epithelial subset showing expression of a signature derived from all genes encoding proteins found in the EpCAM⁺ cell proteome. **(G)** UMAP displaying expression of a refined signature representing overlapping genes from the tissue secretome and EpCAM⁺ cell proteome. **(H)** Bar plot quantifying the number of secreted molecules from the tissue secretome detected across epithelial subclusters. **(I)** Electron microscopy (EM) image of an FT organoid showing secretory granules along the apical surface of multiciliated and secretory cells. Annotations: C = cilia; MV = microvilli; S = secretory granules. Scale bar, 10 µm. **(J)** Table listing the top 20 differentially expressed secretory molecules from each epithelial subcluster. Color shading reflects the presence of each gene’s protein product in the proteome and/or secretome datasets shown in panel **B**, indicating relative support across modalities. ***See Supplementary*** Figure 6 ***for additional related data*.**

We first assembled a reference list of secreted molecules by integrating annotated molecules from public databases, including the Human Protein Atlas and UniProt. To enrich for FT-specific factors, we incorporated proteomic data from our FT tissue and 2D culture secretomes, as well as published datasets from FT lavage fluid ^54^ and PAX8⁺ 2D cultures ^55^. This list was further refined using proteins detected in the EpCAM⁺ epithelial proteome, 3D organoid proteome, and publicly available FT tissue proteomes ^56^. We then mapped these secreted molecules onto epithelial subclusters from our integrated single-cell atlas (**Figure 3D**), enabling resolution of both broad and subcluster-specific secretory programs and their putative biological roles (**Figure 5B**).

To identify proteins secreted specifically by FT epithelial cells, the primary source of the tubal secretome, we first validated the biological relevance of our *in-vitro* tissue secretome by comparing it to *in-vivo* FT lavage fluid analyzed by mass spectrometry. Notably, 84% of proteins detected in lavage fluid were also present in the tissue secretome, supporting its use as a proxy for the native FT secretory environment (**Figure 5C**). We then identified 1,464 proteins shared between the tissue secretome and the EpCAM⁺ epithelial cell proteome, which represents epithelial-derived secreted factors (**Figure 5D**). Comparison of these epithelial-derived secreted proteins with our P0 2D culture secretome revealed 72% overlap, suggesting that early-passage epithelial cultures recapitulate key aspects of FT secretory output (**Figure 5E**).

We next evaluated the representation of epithelial subclusters in the integrated scRNA-Seq dataset within the EpCAM⁺ epithelial cell population. Application of a total proteome signature derived from FT EpCAM⁺ cells to the integrated epithelial dataset revealed broad representation across most clusters, including both secretory and multiciliated lineages (**Figure 5F**). To assess which clusters contribute most to the FT secretome, we applied a signature for epithelial-derived secreted proteins (overlapping proteins in **Figure 5D**) to the integrated epithelial subset. This signature showed the strongest expression in a subset of secretory clusters, though multiciliated clusters also displayed notable expression (**Figure 5G; Supplementary Figure S6A)**. To confirm these findings, we quantified the number of secreted molecules from each epithelial cluster that were also detected in the FT tissue secretome. All clusters were capable of contributing to secretion, and some multiciliated clusters showed greater representation than other secretory clusters (**Figure 5H**). Supporting this, electron microscopy of FT organoids revealed putative apical secretory granules in both multiciliated and secretory cells, further indicating secretory capacity in both lineages (**Figure 5I**).

To explore lineage-specific secretion patterns, we examined the top 20 secreted molecules from each epithelial cluster and annotated their presence across 9 in-house and public proteomic datasets (**Figure 5J**). Notably, cluster 5 is enriched for secreted extracellular matrix (ECM) components such as *COL6A2*, *COL6A3*, *EMILIN1*, *NID1*, *FBN1*, and *SPARC*, suggesting a prominent role in tissue homeostasis and remodeling. Secretory cluster 11 is distinguished by secretion of *OVGP1*, a glycoprotein essential for sperm-oocyte binding;^15^ *SERPINA5*, a serine protease inhibitor that modulates sperm motility;^57^ and *CRISP3*, a glycoprotein implicated in early embryo development.^58^ Among other secretory populations, cluster 9 expresses *LIF*, a cytokine critical for implantation and immune regulation,^59^ alongside several chemokines (*CXCL6*, *CXCL1*, *CCL20*), while cluster 10 secretes *RSPO1*, an agonist of the WNT signaling pathway. In contrast, multiciliated cluster 1 shows enrichment of secreted factors involved in the oxidative stress response, including *TXN* and *GSTA1*. Cluster 0, also multiciliated, appears to secrete molecular components of the ciliary rootlet (*CROCC, CROCC2*) and microtubules (*MAP1A, MAPRE3, SYNE1*), possibly through ciliary extracellular vesicles.^60^ This analysis demonstrated distinct secretion profiles among clusters, suggesting specialized functional roles linked to their biological and reproductive context.

### Subpopulation-Specific Secretory Signatures Link Rare Bi-lineage Tubal Epithelial Cells to Mesenchymal-Like High-Grade Serous Ovarian Cancer

To investigate potential cancer associations between FT epithelial subclusters and HGSOC subtypes, we compared the top 20 secreted molecules from each epithelial cluster to bulk RNA-seq data from 307 HGSOC tumors in The Cancer Genome Atlas (TCGA), which were stratified into the four transcriptional subtypes: differentiated, immunoreactive, mesenchymal, and proliferative.^61^ This revealed significant transcriptional associations between two epithelial clusters and HGSOC subtypes— specifically, cluster 5 with the mesenchymal subtype and cluster 2 with the immunoreactive subtype (**Figure 6A; Supplementary Figure S7A-B)**. To validate these associations at the protein level, we performed a parallel analysis using proteomic profiles from 54 HGSOC tumors from the Clinical Proteomic Tumor Analysis Consortium (CPTAC).^62^ Consistent with the transcriptomic analysis, cluster 5’s secreted protein signature was significantly enriched in tumors of the mesenchymal subtype. In contrast, cluster 2 showed weaker proteomic correlation with the immunoreactive subtype (**Figure 6B; Supplementary Figure S7C-D)**. To further support the mesenchymal molecular profile of cluster 5, we compared its top 50 secreted molecules to microarray data from epithelial and stromal regions of HGSOC tumors, isolated by laser-capture microdissection in Tothill et al.^63^ This analysis revealed a stronger association between cluster 5 and the tumor stroma than the tumor epithelium, based on the expression patterns of these secreted molecules **(Supplementary Figure S7E)**. Given that the mesenchymal subtype is associated with the poorest prognosis among HGSOC subtypes,^64^ and that cluster 5 exhibited the strongest and most consistent transcriptomic and proteomic associations with this subtype, we focused subsequent analyses on characterizing this cellular population in greater depth.

**Figure 6.**
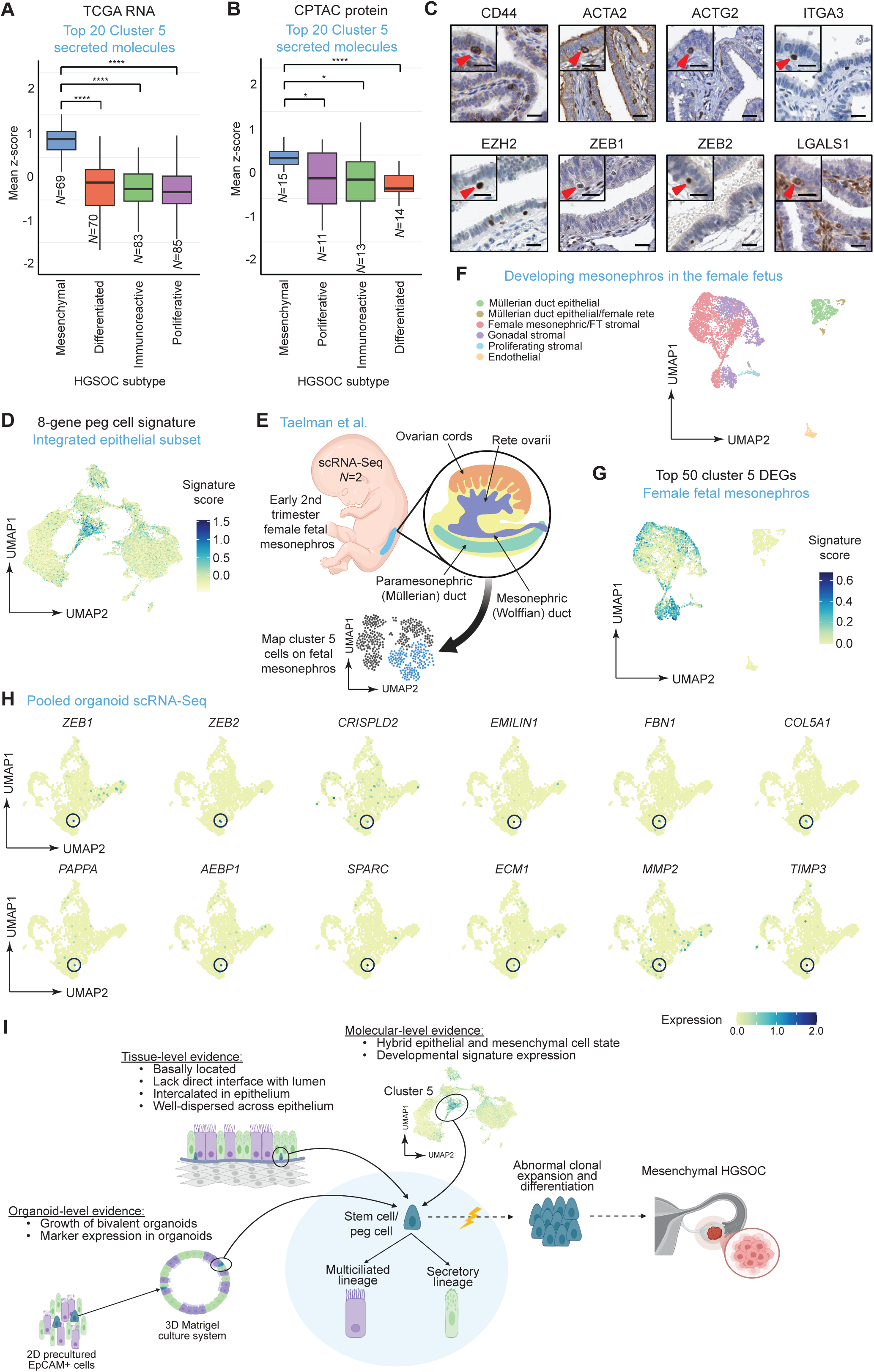
Subpopulation-specific secretory signatures link rare mesenchyme-like bi-lineage tubal ‘peg’ cells to mesenchymal-like HGSOC and fetal mesonephric stromal programs. **(A)** Boxplot showing the mean z-score of the top 20 secreted molecules from epithelial subcluster 5 compared to RNA-seq data from 307 patients across the four TCGA-defined high-grade serous ovarian cancer (HGSOC) subtypes. **(B)** Boxplot comparing the same 20 secreted molecules from subcluster 5 with CPTAC proteomic data from 54 HGSOC patients across the four subtypes. **(C)** Immunohistochemical staining images from the Human Protein Atlas showing expression of mesenchymal markers enriched in subcluster 5, highlighting their localization to basally positioned peg-like cells (Source: Human Protein Atlas, https://www.proteinatlas.org/). Large image scale bars, 25 µm. Inset scale bars, 20 µm. **(D)** Expression of an 8-gene peg cell signature (derived from panel **C**) visualized on the UMAP of the integrated epithelial subset. **(E)** Experimental workflow for assessing transcriptional similarity between subcluster 5 and a publicly available single-cell RNA-seq dataset of human female fetal mesonephric tissue. **(F)** UMAP shows annotated cell types in the fetal mesonephric dataset. **(G)** Application of a gene signature comprising the top 50 differentially expressed genes from subcluster 5 to the fetal mesonephric UMAP. **(H)** Feature plots showing expression of top secreted molecules and peg markers from subcluster 5 projected onto a pooled organoid scRNA-seq UMAP (*N*=2). **(I)** Schematic summary integrating organoid, tissue, and molecular-level evidence supporting the existence of a peg-like stem cell population in subcluster 5 with potential relevance as a cell of origin for mesenchymal HGSOC. ***N***=patients; **n**= specimens (applies to all panels). Significance was determined by two-sample t test with Benjamini-Hochberg FDR correction; *, *p* < 0.05; ****, *p* < 0.0001. ***See Supplementary*** Figure 7 ***for additional related data*.**

### Mesenchymal-Like Features Define the HGSOC-Associated Epithelial Subpopulation

We first confirmed that cluster 5 maintains an epithelial identity with expression of mesenchymal features. All 16 epithelial clusters in our integrated scRNA-Seq dataset expressed canonical epithelial markers including *EpCAM*, cytokeratins, and claudin-family genes **(Supplementary Figure S7F)**. In contrast, cluster 5 uniquely expressed high levels of mesenchymal and ECM-associated genes such as *CRISPLD2*, *NID1*, *EMILIN1*, *COL6A3*, and *FBN1* **(Supplementary Figure S7G)**. To exclude the possibility that cluster 5 reflects stromal contamination, we compared expression of fibroblast, myofibroblast, and smooth muscle markers across cluster 5, the three stromal clusters from the integrated dataset, and all other epithelial subclusters. While stromal clusters exhibited robust expression of these markers, only cluster 5 showed low expression, with all other epithelial clusters showing minimal to no expression **(Supplementary Figure S7H)**. Further, cluster 5 cells displayed a unique gene expression profile enriched for ECM regulators, including select laminins and collagens, suggesting a distinct basement membrane-modifying capacity **(Supplementary Figure S7I)**.

### Mesenchyme-like Epithelial Subpopulation (Cluster 5) Identifies Rare Basal ‘Peg’ Cells

To spatially localize cluster 5 cells within FT tissue, we used immunohistochemistry data from the Human Protein Atlas to assess the expression of secreted and mesenchymal subtype-associated markers. This analysis identified a consistent panel of eight proteins (CD44, ACTA2, ACTG2, ITGA3, EZH2, ZEB1, ZEB2, and LGALS1) that are present in a rare, basally positioned subset of cells intercalated among the epithelial layer, lacking direct luminal contact and well-dispersed throughout the tissue (**Figure 6C**). This rare population resembles previously described EpCAM^+^/CD44^+^/ITGA6^+^ “peg” or “intercalated” cells.^10,11,41^ To test whether cluster 5 corresponds to this histologically defined population, we generated an 8-gene signature from these proteins and applied it to the epithelial subset of the integrated dataset. Signature scoring revealed selective enrichment in cluster 5, supporting the hypothesis that cluster 5 represents peg-like cells with secretory and mesenchymal features (**Figure 6D**).

### Rare Basal ‘Peg’ Cells Maintained *Ex-Vivo* Exhibit Transcriptional Parallels with Fetal Multipotent Mesonephric Stromal Cells

To investigate whether the transcriptional identity of cluster 5 cells reflects a developmental origin, we compared their molecular signature to cell populations present in the female fetal mesonephros, a transient embryonic structure that gives rise to the paramesonephric (Müllerian) ducts and components of the ovarian stroma (**Figure 6E**). Scoring single-cell transcriptomes from 16–18-week human fetal mesonephros^65^ with the top 50 differentially expressed genes from cluster 5 revealed strong enrichment within mesonephric and fallopian tube stromal cell populations, as well as early gonadal stromal compartments (**Figure 6F-G**). These findings suggest that cluster 5 cells share key molecular features with multipotent progenitor populations arising during mesonephric and Müllerian development.

We next asked whether this cell population is sustained *in-vitro*. Analysis of pooled organoid scRNA-seq data revealed a small but transcriptionally distinct epithelial subset (∼0.1% of cells) expressing the top secreted genes characteristic of cluster 5. As the cluster 5 cell population displays stem-like features, its low frequency yet consistent presence in organoids suggests it may be enriched for organoid-initiating cells (**Figure 6H**).

Taken together, these results define a rare basal epithelial subpopulation with mesenchymal and secretory features, transcriptionally aligned with fetal mesonephric progenitors and preserved *ex-vivo*. The presence of this population in both primary tissue and culture models nominates it as a candidate cell type for studying epithelial plasticity and subtype-specific transformation in the fallopian tube epithelium (**Figure 6I**).

## Discussion

Here, we describe the establishment of the largest clinically and genetically annotated FT organoid biobank to date—a scalable, renewable platform for dissecting FT epithelial biology at single-cell resolution and modeling its dysregulation in cancer. At the time of submission, the biobank comprises tissue and Tao brushing samples from 202 living and deceased women, encompassing a broad spectrum of ages, germline mutation statuses, reproductive histories, and cancer risk profiles. By integrating organoid technology with high-dimensional single-cell and multi-omic profiling, this platform offers unprecedented insight into the cellular hierarchies and transcriptional programs governing FT homeostasis and provides a tractable system for modeling early events in HGSOC pathogenesis.

To resolve the molecular architecture of the FT epithelium, we implemented an integrative multi-omic framework spanning primary tissues and matched organoid models. By combining single-cell and single-nucleus transcriptomic profiling with epithelial-specific proteomic and secretome analyses, we generated the most comprehensive FT cell atlas to date. This approach delineated high-resolution trajectories of multiciliated and secretory lineage commitment and revealed transcriptionally distinct epithelial subpopulations with potential functional specialization. Unlike traditional lineage markers prone to intrapatient and intrastudy variability, such as *PAX8* and *FOXJ1*, we derived and validated robust six-gene lineage signatures that were highly specific and reproducible across scRNA-seq, snMultiome, and bulk modalities, thereby improving the ability to accurately identify and differentiate multiciliated and secretory lineages. Applying these signatures, we show that our culture system consistently generates bi-lineage organoids, though the lineage composition is strongly biased toward the secretory compartment which is further supported by ultrastructural characterization using electron microscopy. While several studies have identified pathways and media modifications that promote ciliogenesis,^20,37^ our culture conditions do not incorporate all of these factors, likely contributing to the reduced representation of multiciliated cells *in-vitro*. Interestingly, the secretory nature of the organoids more closely resembles the secretory-like epithelial cells that comprise early precursor lesions such as p53 signatures and STICs, increasing their relevance as a model for transformation-prone epithelial states.^4–6^ Ultimately, these signatures provide a scalable and accurate framework for epithelial lineage classification in both physiologic and pathologic contexts, enabling future efforts to map early transformation events in HGSOC.

Leveraging chromatin accessibility and gene regulatory network inference allowed us to extend beyond gene expression to define the regulatory transcriptional programs underpinning these epithelial identities. Multiciliated cells were shown to be regulated by *RFX* and *FOX* family TFs, consistent with their roles in ciliogenesis, while secretory cells exhibited a more diverse regulatory landscape involving *SOX*, *TEAD*, and *NFI* family members. Notably, *PAX8*, commonly used as a defining secretory marker,^66–68^ emerged not as a standalone regulator, but as part of a broader previously undescribed transcriptional network. Although PAX8 and SOX17 have been shown to physically interact in normal FT epithelium and HGSOC, we show that neither factor transcriptionally regulates the other’s expression.^69^ Altogether, this highlights the complexity of secretory lineage specification and the importance of complex TF activity in maintaining epithelial fate.

Beyond canonical lineage distinctions, our data reveal functional heterogeneity in secretory activity across transcriptionally defined epithelial subclusters. Integrated proteomic and transcriptomic analyses demonstrate that both multiciliated and secretory lineages contribute to the FT secretome, with discrete subpopulations exhibiting distinct secretory profiles. Importantly, this study is the first to link these epithelial subclusters to putative reproductive functions through the expression of well-characterized reproductive molecules, offering new insights into FT physiology and functional specialization. These specialized sets of secreted molecules suggest a spatial and functional compartmentalization of epithelial signaling, extending beyond traditional lineage boundaries. This distributed secretory architecture likely underpins key physiological functions, ranging from reproductive tract conditioning and immune modulation to epithelial barrier maintenance, and may inform how early alterations in secretory states contribute to HGSOC risk. Further work is needed to define how these compartmentalized secretory programs are dynamically regulated by hormonal cues and menstrual cycling.

Among the most compelling discoveries of this study is the identification of a rare epithelial subpopulation, cluster 5, that exhibits transcriptomic and proteomic similarity with the mesenchymal subtype of HGSOC, the most aggressive and therapy-resistant form of the disease.^64^ Cluster 5 cells co-express hallmark epithelial genes alongside mesenchymal and extracellular matrix remodeling factors, consistent with a stable hybrid epithelial– mesenchymal phenotype, which we detect across both primary tissue and organoid models. Cluster 5 cells show strong correlation with rare epithelial subsets described by Ulrich et al.,^18^ including the non-ciliated secretory epithelial clusters NCSE 2-1 and NCSE 2-2 and the ciliated epithelial cluster CE 1-4, all of which express combinations of mesenchymal-associated genes such as *ACTA2, COL1A1, MMP2, PRRX1, DCN, FN1, TIMP3, LGALS1, SPARC*, and *TAGLN*. Additionally, in their study, some of these cells were validated using immunofluorescence for EPCAM/ACTA2 double-positive cells in fimbria, ampulla, and isthmus tissues. Furthermore, cluster 5 cells strongly map to unclassified cluster 1 cells from Dinh et al.,^34^ which were enriched in early timepoints of pseudotime analysis thus suggesting their potential role as progenitors or cells early in differentiation. The unclassified cluster 1 cells were also mapped to TCGA HGSOC RNA-Seq profiles where they were enriched in ∼52% of tumors, with the highest percentage of the mesenchymal subtype.^34^ In our study, we show a specific and highly significant enrichment of secretory molecules from cluster 5 cells in the mesenchymal subtype of HGSOC. Thus, secreted molecules from cluster 5 warrant future evaluation as potential biomarkers for disease initiation, subtype stratification, and progression monitoring.

Further, the enrichment of fetal mesonephric stromal signatures within cluster 5 suggests developmental continuity, implicating a peg-like progenitor population that persists from fetal stages into adulthood. Their basal localization, scarcity, and molecular identity align with historical descriptions of “peg” cells ^10,11,41^, yet this study provides the first integrated single-cell and proteomic framework to resolve their phenotype. These findings nominate cluster 5 as a multipotent stem-like epithelial subset with potential roles in maintaining epithelial homeostasis and serving as a candidate cell-of-origin for the mesenchymal subset of HGSOC.

The identification of cluster 5 gains further relevance in the context of aging which is a key risk factor for transformation. While aging is associated with reduced epithelial content and a decline in multiciliated cells,^52^ organoid-forming capacity remains stable, suggesting that progenitor function is retained. These shifts likely reflect altered differentiation potential or microenvironmental influences, rather than stem cell exhaustion, underscoring the need to disentangle intrinsic and extrinsic drivers of epithelial plasticity with age. In support of this, we identified age- and menopause-associated changes in epithelial composition, including a progressive decline in multiciliated populations (clusters 0 and 1) and a selective expansion of one secretory subpopulation (cluster 4). These changes suggest a shift in the physiological distribution of epithelial subtypes with age which may create a cellular context more susceptible to oncogenic transformation.

Together, this work provides a clinically annotated, multi-omic platform for modeling FT epithelial dynamics and identifying candidate cells of origin for HGSOC. The diversity of samples enables systematic interrogation of how patient-specific variables, such as *BRCA1/2* and *TP53* mutations, hormonal exposures, and prior malignancy, shape epithelial composition, lineage allocation, and transformation potential. The integration of patient-derived organoids with single-cell and proteomic profiling offers a scalable framework to enhance our knowledge of lineage commitment, transformation risk, and subtype-specific vulnerability. Moving forward, the isolation of distinct epithelial subsets, including cluster 5, will be essential to establish their contributions to epithelial homeostasis and lineage commitment. By aligning our biobank with emerging spatial maps of the FT in precancerous states and with new insights into chronic inflammatory and oncogenic stromal interactions,^70^ we will be positioned to trace the earliest cellular perturbations at single-cell resolution and define the pathways by which normal progenitors are recruited into malignant transformation.

### Limitations of the study

While our study centers on the epithelial origin of HGSOC, other cell types may shape its origins,^71^ and their roles await future spatially resolved investigation.

## Methods

### Donor eligibility and informed consent for specimen collection

The fallopian tube (FT) organoid biobank is an opt-in research resource for which written informed consent is obtained from eligible participants. All human subject procedures are conducted in accordance with the Declaration of Helsinki and are approved by the Mayo Clinic Institutional Review Board (IRB# 18-001967). Women aged ≥18 years undergoing salpingectomy or other gynecologic surgeries involving the FT are eligible to participate, provided they are capable of understanding and signing an informed consent document. Exclusion criteria include any history of psychiatric or neurological conditions that may impair comprehension or capacity to consent. Consent may be obtained in person or remotely via secure electronic platforms. Autopsy cases are identified and referred by the Mayo Clinic Tissue Registry and Autopsy Group (TRAG); for these donors, tissue is collected post-mortem under the approved IRB protocol without the requirement for prospective consent. Enrollment for the biobank started at Mayo Clinic Rochester in 2018, with active ongoing collections.

Patient recruitment is conducted by trained study staff and coordinators, who review clinical schedules up to one month in advance. When a potentially eligible patient is identified, the attending surgical team is informed, and the patient is contacted by a study coordinator. If the patient agrees to participate, consent is obtained prior to surgery. Participants authorize the use of tissue or Tao brushing samples for organoid generation, as well as storage of clinical data and biospecimens for future research. For all living participants (high- or low-risk), excess surgical specimens not required for clinical pathology are collected from the frozen section laboratory and transported to the research laboratory. Archived clinical tissue from other body sites may also be retrieved from the Mayo Tissue Registry for participants who have provided consent. Autopsy-derived tissue is limited to fresh FT samples obtained by TRAG and transferred directly to the research team for biobanking.

Biospecimens, including organoids, cell suspensions, and tissue fragments, are tracked in the Mayo Research Laboratory Information Management System (RLIMS), and will be cryopreserved for long-term storage or used for current experiments. RLIMS is used to assign and track specimen IDs, protocol numbers, and banking codes. All identifiers are securely coded and linked to patient information via protected research drives; no names or medical record numbers are accessible in shared or distributed materials.

Participants may withdraw from the biobank at any time. In such cases, they are asked whether previously collected data or biospecimens may continue to be used. If declined, all identifiable materials are destroyed and excluded from future research. Risks to participants are minimal and limited to the rare possibility of unintended disclosure of personal information. All specimens and data are handled in accordance with institutional privacy and confidentiality protocols.

All requests for specimens and data will be thoroughly reviewed in accordance with Mayo Clinic policy and alignment with collaborative goals to advance knowledge in FT epithelial and cancer stem cell biology. For further details, please email the corresponding authors.

### Viable FT specimen processing and annotated biobanking

FT epithelial cells were isolated and expanded using our previously published protocol optimized for both tissue and Tao brushing samples (Cook Medical, G17023).^40^ Briefly, fresh surgical tissue or Tao brushing specimens were transported to the laboratory in cold transport medium and processed within 2–3 hours. Tissues were enzymatically dissociated with Collagenase I (EMB Millipore, SCR103, 3301502) and mechanically disaggregated via scraping with a scalpel to release epithelial cells, followed by short-term 2D expansion in defined epithelial medium. Length of enzymatic dissociation (45 minutes or overnight) was tested and optimized. For Tao brushing samples, red blood cell lysis and 2D culture were incorporated to improve epithelial enrichment prior to 3D Matrigel (R&D Systems, 3533-005-02) embedding.

### Fallopian tube 2D culture

Isolated single cells from tissue or Tao brushing processing were cultured in 2D, embedded directly in Matrigel for 3D culture or precultured in 2D prior to 3D culture. Briefly, for 2D culture, cells were seeded in 60 mm or 10 cm culture dishes with 3 or 8 mL of FT cell culture medium respectively: Advanced DMEM/F12 (Gibco, 1998031) supplemented with 12 mM HEPES (Gibco, 15630-080), 1X GlutaMAX (Gibco, A12860-01), 10 ng/mL EGF (Sigma-Aldrich, E9644), 9 uM ROCK inhibitor (Y-27632, EMD Millipore, 2975574), 5% bovine serum (Gibco, 26170-043) and 1X Penicillin-streptomycin (Gibco, 15140-122) or 1X Anti-Anti (Gibco, 15240062). Cultures were maintained until confluency approached 70-80%, after which they were subcultured, embedded in Matrigel (R&D Systems, 3533-005-02) or cryopreserved in liquid nitrogen.

### Flow cytometry and fluorescence-activated cell sorting (FACS)

FT cells from fresh tissues, 2D culture expanded cells at different passages, or cultured 3D-matrigel organoids were analyzed and/or sorted by flow cytometry/ FACS. For fresh tissues, a single cell suspension was generated using the above protocol for tissue dissociation. For 2D cultures, cells were trypsinized with TrypLE™ Express (Gibco, 12605-010) for 5 minutes and collected via centrifugation at 300 x g at 4°C for 5 minutes. To isolate single cells from organoids, the organoid-matrigel domes were resuspended in 100 μL of Cell Recovery Solution (Corning, 354253) and incubated at 4°C for 30 minutes. The solution was then manually pipetted to disrupt the organoids followed by centrifugation at 2500 rpm for 5 minutes. The supernatant was removed and 500 uL of TrypLE™ Express (Gibco, 12605-010) was added. The solution was mixed by pipetting up and down and incubated at 37°C for 5 minutes. 1 mL of HBSS (Gibco, 14175-095) supplemented with 2% FBS (Sigma-Aldrich, F1051) was then added prior to subsequent centrifugation to yield a single cell pellet.

The cells were stained with the following anti-human fluorochrome-conjugated antibodies: 1:50 EpCAM PE (BioLegend, 324206,Clone 9C4), 1:50 Integrin α6 APC (R&D Systems, FAB13501A), 1:50 CD45 Pacific Blue™ (BioLegend, 304029, Clone HI30), 1:50 CD31 Pacific Blue™ (BioLegend, 303114, Clone WM59) and 1:200 DAPI (4′,6-diamidino-2-phenylindole; Thermo Fisher Scientific, Cat. #D1306). Stained cells were analyzed using FACSMelody cell sorter/analyzer (BD Biosciences). For sorting, hematopoietic (CD45+), endothelial (CD31+), and dead (DAPI+) cells were removed leaving lineage-negative (Lin-) cells. From Lin-cells, viable subsets of EpCAM+/-cell populations were gated and sorted at ≥98% purity. The FCS data files were analyzed using FlowJo data analysis software package (version 10.8.1). Approximately 25,000–1,500,000 cells were sorted or analyzed in an experiment.

### 3D Matrigel culture and organoid-initiating cell (OIC) activity assay

Fresh P0 and FACS-sorted FT cells were optimally precultured in 2D for 1-2 weeks (some up to 4 weeks) until they reached 70-80% confluence. Cells were cultured on either untreated or collagen treated (Stem Cell Technologies, 04902) plates and maintained in incubators with either 5 or 20% oxygen. Twenty-five thousand cells were plated in 96 well plates as 20 uL Matrigel domes (R&D Systems, 3533-005-02). To allow the matrigel to solidify, the domes were incubated at 37°C for 15-30 minutes. Subsequently, 200 uL of FT organoid media containing Advanced DMEM/F12 (Gibco, 1998031) supplemented with 12 mM HEPES (Gibco, 15630-080), 1x GlutaMAX (Gibco, A12860-01), 10 ng/mL EGF (Sigma, E9644), 9 uM ROCK inhibitor (Y-27632, EMD Millipore, 2975574), 25% WNT3a conditioned media,^40^ 25% RSPO1 conditioned media,^40^ 1x B-27 (Gibco, 17504044), 1x N2, 100 ng/mL Noggin (PEPROTECH, 120-10C-20ug), 100 ng/mL FGF10 (PEPROTECH, 100-26-25ug), 1 uM Nicotinamide (Sigma-Aldrich, N0636) and 0.5 uM TGFBR1 Kinase Inhibitor IV (SB431542, EMD Millipore, 909910-43-6) were added to each well. Media changes were completed 3 times per week. At experimental timepoints or endpoints, organoids per well were counted under a brightfield microscope and the frequency of 3D organoid initiating cells (OIC) was calculated by dividing the number of organoids formed by the number of cells plated.

For organoid cryopreservation, organoid cultures were washed with 4°C prechilled Cell Recovery Solution (Corning, 354253), the Matrigel was scraped from the bottom of the plate with a pipette and transferred to a 15 mL Falcon tube with cold buffer. The organoids were centrifuged at 300 x g at 4°C for 5 minutes, supernatant was removed, and they were then resuspended in cryomedium (50% Advanced DMEM/F12, 44% bovine serum, 6% DMSO (Sigma, D4545)). To revive organoids, the cryovials were rapidly thawed in the 37°C water bath and washed in PBS (Corning, 21-040-CV) prior to being plated in 20-50 uL of Matrigel and covered with 200 uL organoid growth medium.

### Immunohistochemistry

For immunohistochemistry, paraffin-embedded organoid sections were deparaffinized and subjected to antigen retrieval using Antigen Unmasking Solution (Vector Laboratories, Burlingame, CA) according to the manufacturer’s instructions. Sections were subsequently stained with hematoxylin and eosin, and all slides were imaged using the Cytation 5 Imaging System (BioTek Instruments, Agilent Technologies).

### Electron microscopy

Serial block-face scanning electron microscopy (SBFSEM) was performed using a double osmium (OTO) protocol previously described.^72^ Briefly, dissected FT tissue and cultured FT organoids were fixed by immersion in 2% glutaraldehyde + 2% paraformaldehyde in 0.1 M cacodylate buffer containing 2 mM calcium chloride. Organoids were then suspended in 2% agar, solidified, and further processed same as tissue. After fixation, the sample was rinsed in 0.1 M cacodylate buffer and placed into 2% osmium tetroxide + 1.5% potassium ferracyanide in 0.1 M cacodylate, washed with nH_2_O, incubated at 50°C in 1% thiocarbohydrazide, incubated again in 2% osmium tetroxide in nH_2_O, rinsed in nH_2_O and placed in 2% uranyl acetate for 12 hours. The next day the sample was rinsed again in nH_2_O, incubated with Walton’s lead aspartate, dehydrated through an ethanol series, and embedded in Embed 812 resin (EMS, Hatfield, PA). To prepare the embedded sample for serial sectioning, a 1.0 mm^3^ area was trimmed of any excess resin and mounted to an 8 mm aluminum stub using silver epoxy Epo-Tek^®^ H20E (EMS, Hatfield, PA). The mounted sample was then carefully trimmed into a smaller 0.5 mm^3^ tower using a Diatome^®^ trim 90 diamond trimming tool (EMS, Hatfield, PA) and vacuum sputter-coated with gold to help dissipate charge. Serial sectioning and imaging of the sample was performed using VolumeScope 2 SEM^TM^ (Thermo Fisher Scientific, Waltham, MA). Imaging was performed under low vacuum mode to suppress charging artifacts, with a starting energy of 3.0 keV and beam current of 0.10 nA. Approximately 300 images were acquired at 10 nm x, y resolution with a z step of 75 nm (organoid) or 50 nm (tissue). Image registration and volume rendering were performed using Amira software (Thermo Fisher Scientific) with volume renderings colorized in Photoshop (Adobe).

### DNA isolation from FT FFPE blocks and snap frozen tissue

Genomic DNA was isolated from FFPE tissue blocks using the QIAamp® DNA FFPE Tissue Kit (Qiagen, 56404) and Deparaffinization Solution (Qiagen, 19093) according to the manufacturer’s protocol. Briefly, 5–10 µm sections were cut from FFPE blocks without trimming excess paraffin, as samples were obtained from the clinical tissue registry. Because of the excess paraffin, maximum volumes of Deparaffinization Solution were used. Tissue sections were incubated with Deparaffinization Solution at 56°C for 3 minutes, followed by lysis in Buffer ATL and Proteinase K. After a 1-hour digestion at 56°C, samples were further incubated at 90°C for 1 hour to partially reverse formaldehyde crosslinking. Following lysis, samples were subjected to ethanol precipitation and transferred to QIAamp MinElute columns. DNA was purified using standard binding, washing, and elution steps per the QIAamp protocol, with final elution in Buffer ATE.

Genomic DNA was isolated from snap frozen tissues using the MasterPure Complete DNA and RNA Purification kit (Biosearch Technologies, MC85200) according to the manufacturer’s protocol for tissue samples. Briefly, 1–5 mg of tissue was homogenized or ground in liquid nitrogen, then lysed in Tissue and Cell Lysis Solution containing Proteinase K and incubated at 65 °C for 15 minutes with intermittent vortexing. After lysis, samples were cooled on ice, treated with RNase A, and subjected to protein precipitation using MPC Protein Precipitation Reagent. DNA was pelleted with isopropanol, washed with 70% ethanol, and resuspended in TE Buffer (Invitrogen, 12090015). DNA concentrations and quality were quantified using an IMPLEN NanoPhotometer®.

### ScRNA-Seq of FT single cells and organoids

Dissociated fresh cells and organoids were pelleted, resuspended in PBS (Corning, 21-040-CV), and stained with Trypan Blue Solution 0.4% (Gibco, 15250-061) and counted using a hemocytometer. The volume of cell suspension was adjusted to 400-1000 cells/ul in PBS. Cells were then processed on a 10x Genomics Chromium Controller using the Chromium Next GEM Single Cell 3′ reagents kit v3.1 (Dual Index) (10x Genomics, PN-1000268) following the manufacturer’s instructions. In brief, live cells were loaded onto the Chromium controller to recover approximately 10,000 cells for library preparation and sequencing. Gel beads were prepared following the manufacturer’s instructions. Subsequently, oil partitions of single-cell and oligo-coated gel beads were captured, and reverse transcription was performed, resulting in cDNA tagged with a cell barcode and unique molecular index (UMI). Next, GEMs were broken, and cDNA was amplified and quantified using an Agilent Bioanalyzer High Sensitivity chip (Agilent Technologies).

To prepare the final libraries, amplified cDNA was enzymatically fragmented, end-repaired, and polyA tagged. Fragments were then size-selected using SPRIselect magnetic beads (Beckman Coulter, B23317). Next, Illumina sequencing adapters were ligated to the size-selected fragments and cleaned up using SPRIselect magnetic beads. Finally, sample indices were selected and amplified, followed by a double-sided size selection using SPRIselect magnetic beads. Final library quality was assessed using an Agilent Bioanalyzer High Sensitivity chip. The libraries were then sequenced as paired end reads (PE150) on Novaseq platform (Illumina).

### Nuclei preparation for snMultiome sequencing

Fresh fallopian tube tissues were processed for single-nucleus ATAC and RNA sequencing using the 10x Genomics Single Cell Multiome protocol for complex tissues as previously described (10x Genomics, Cat#PN-1000283).^73^ For nuclei extraction, a small piece of tissue was homogenized on ice in 300 µL NP-40 Lysis Buffer using a pellet pestle, followed by the addition of 1 mL NP-40 Lysis Buffer and incubation on ice for 5– 30 minutes. Lysates were filtered through a 70 µm strainer, and nuclei were pelleted at 500 × g for 5 minutes at 4°C. The pellet was resuspended in 100 µL of 0.1× Lysis Buffer for 2 minutes, washed with 1 mL Wash Buffer, and centrifuged again. Final nuclei were resuspended in Diluted Nuclei Buffer and counted by PI staining using a Cellometer K2. Approximately 5,000 nuclei were targeted per sample.

Nuclei (1,000–8,000/sample) were tagmented with Tn5 transposase and processed using the Chromium platform for droplet partitioning, barcoding, and cDNA preamplification. ATAC and gene expression libraries were prepared per 10x Genomics protocol, quantified with Qubit High Sensitivity assays, and assessed via Fragment Analyzer. Libraries were sequenced for paired-end 50 bp on Illumina HiSeq 4000, PE50 on NextSeq 2000, and 100PE on NovaSeq X 1.5B systems. Demultiplexing and alignment were performed using GRCh38 (hg38) as the reference genome.

### scRNA-Seq data processing and quality control

Raw sequencing data from fresh tissue and organoids were processed using Cell Ranger v6.1.2 (10x Genomics) against the GRCh38 reference genome (v2020-A) to generate filtered gene-barcode matrices for each sample. Subsequent analysis was performed in R (v4.2.2) using Seurat v4.4.0.^74^ Samples were imported using *Read10X* and converted into a Seurat object with *CreateSeuratObject* using a minimum of 200 detected features and 3 cells. Cells were filtered out if they fell outside the following QC thresholds: 200– 7,000 detected features, <20% mitochondrial content, >1% ribosomal content, and <0.5% hemoglobin content. Following filtering, gene expression was normalized using log-normalization (*NormalizeData*). Highly variable genes were identified with *FindVariableFeatures* using the “vst” method (N=2,000). Datasets were integrated using anchor-based integration via *FindIntegrationAnchors* and *IntegrateData*. Dimensionality reduction was performed with *RunPCA.* Graph-based clustering was performed using *FindNeighbors* and *FindClusters* with a resolution of 0.5 and samples were embedded using *RunUMAP*. Cell type identification was based on canonical marker gene expression and differential expression testing. Marker genes were identified using *FindAllMarkers* with a Wilcoxon Rank Sum test using min.pct=0.25, and logfc.threshold=0.25.

### snMultiome data processing and quality control

Single-nucleus multiome (snATAC + snRNA-seq) data were processed using a previously published pipeline.^73,75^ Sequencing reads were aligned to the GRCh38 reference genome (10x Genomics v2020-A-2.0.0) using Cell Ranger ARC v2.0.0. Following quality control, gene expression (GEX) and chromatin accessibility (ATAC) data were analyzed using Seurat^74^ and Signac^76^. GEX matrices were log-normalized, scaled, and reduced using PCA on the top 2,000 variable genes. Uniform manifold approximation and projections (UMAPs) were generated using the top 50 principal components. ATAC peaks were merged across samples (GenomicRanges), filtered by size (20–10,000 bp), and normalized using Term Frequency-Inverse Document Frequency (TF-IDF). Dimensionality reduction was performed via latent semantic indexing (LSI), and UMAPs were computed using LSI components 2–50. Joint modality integration used Seurat’s weighted nearest neighbor (WNN) method, combining GEX PCs 1–50 and ATAC LSI components 2–50 into a shared UMAP. Low quality cells were then filtered out (e.g., >20% mitochondrial reads, <200 genes or peaks, TSS enrichment <1).

### Integration of in-house and public scRNA-Seq and snMultiome sequencing datasets

To generate a comprehensive atlas of human FT epithelial states, we integrated scRNA-seq and single-nucleus multiome (snMultiome) data from both in-house and publicly available datasets. The following studies were included: (1) in-house snMultiome, (2) in-house scRNA-Seq, (3) in-house pooled organoids and external datasets from (4) Dinh et al., (5) Lengyel et al., (6) Weigert et al., (7) Ulrich et al., and (8) Yu et al. Because of variability in public data formats, each dataset was processed into an individual Seurat object prior to integration.^74^ If the available data was in the format of h5 files (Dinh et al.) or barcode matrices (Ulrich et al., Yue et al.), the above scRNA-Seq data processing procedure was used (same as for in-house data). Some studies had data publicly available in Seurat object format (Lengyel et al., Weigert et al.).

The individual Seurat objects were then loaded and labeled with a dataset-specific identifier. Individual objects were normalized independently using *SCTransform* from Seurat and merged using *merge* with cell ID prefixes. The merged object underwent a second round of *SCTransform* normalization to harmonize gene expression scaling across datasets. A set of 3,000 shared variable features was selected for dimensionality reduction. Principal component analysis (PCA) was performed, followed by batch correction using the *RunHarmony* function (Harmony v1.2.3)^77^, with dataset source as the integration variable. Cells were filtered based on quality control criteria: >200 and <7,000 detected genes and <15% mitochondrial reads. Dimensionality reduction, clustering, and visualization were then performed on Harmony embeddings. Nearest neighbor (*FindNeighbors*) graphs were constructed, and clusters were identified with the Louvain algorithm (*FindClusters*, resolution= 0.5). UMAP was computed using the first 25 Harmony dimensions. Integrated data were used for downstream annotation and comparative analyses.

Cell type identities were assigned based on curated marker gene lists representing major FT cell types. Expression patterns of canonical markers were visualized using dot plots to guide manual annotation of clusters. To identify differentially expressed genes across cell states, Seurat’s *FindAllMarkers* function was applied using logistic regression with sample as the testable variable.

### Development of epithelial lineage signatures

Epithelial lineage-specific gene signatures were derived by applying Seurat’s *FindMarkers* function using logistic regression based on samples to compare multiciliated and secretory cell populations, as defined by canonical markers, within both individual datasets and the integrated dataset. Candidate genes were evaluated using *FeaturePlot* to assess expression patterns across the target lineage and all other cell types. Genes exhibiting consistent, robust expression within the lineage of interest and minimal expression in other cell types were retained and compiled into multi-gene signatures. To validate these signatures, *AddModuleScore* was used to calculate lineage-specific scores at the single-cell level in the integrated dataset and in independent fallopian tube datasets, enabling confirmation of lineage specificity and reproducibility across patients and platforms.

### Organoid preparation for bulk RNA-Seq

Organoids were picked up mechanically under the microscope into a sterile 1.5 ml tube, followed by one wash at 350 x g for 10 minutes with cold PBS (Corning, 21-040-CV) to remove residual media and extracellular matrix components. The organoid pellets were then lysed directly in TRIzol™ Reagent (ThermoFisher Scientific, 15596-026) according to manufacturer’s protocol. In brief, total RNA was extracted following phase separation with chloroform and isopropanol precipitation, and RNA pellets were washed with 75% ethanol, air-dried, and resuspended in RNase-free water. RNA integrity and concentration were assessed using IMPLEN NanoPhotometer®, Agilent Bioanalyzer and Qubit fluorometer. Only samples with RNA Integrity Number (RIN) values ≥ 8 were used for downstream processing. Bulk RNA sequencing libraries were prepared and sequenced on the Illumina NovaSeq X Plus platform with paired-end 150 bp reads (PE150). Quality control of raw sequencing data was performed prior to alignment and downstream analysis using FastQC and MultiQC to ensure high-quality reads with minimal adapter contamination and low base-calling error rates.

### Bulk RNA-Seq data processing and quality control

Total RNA from FT organoid samples was sequenced and processed through a standardized pipeline. Quality control on raw reads was performed using FastQC (v0.11.9),^78^ and adapter trimming was conducted using Trim Galore.^79,80^ Post-trimming quality metrics were reassessed to ensure data integrity. Trimmed paired-end reads were aligned to the human reference genome (GRCh38) using STAR (v2.7.9a),^81^ with quantification enabled via the --quantMode TranscriptomeSAM GeneCounts option. Alignment outputs, including BAM files and splice junction tables, were organized into structured directories and indexed using samtools (v1.10).^82^

Gene-level count matrices were generated using *featureCounts* from the Rsubread package,^83^ with in-built hg38 annotations. Downstream analysis was performed in R using *DESeq2* (v1.38.3).^84^ Raw counts were filtered to remove low-expression genes (row sums ≤ 1), normalized using the median-of-ratios method, and transformed using variance-stabilizing transformation (VST) for visualization. Differential expression analysis was conducted using Wald tests and shrinkage estimation (via apeglm)^85^ to identify genes with |log₂ fold change| > 1 and adjusted p-value < 0.05. Quality assessment included PCA, sample clustering, and heatmaps of top variable genes. To derive organoid signature scores from bulk RNA-seq data, gene expression matrices were first filtered to remove housekeeping genes, ribosomal protein genes, and non-epithelial transcripts. Only genes expressed in at least 10 out of 15 samples were retained. The resulting curated gene sets were used to calculate signature scores using the *AddModuleScore* function, based on variance-stabilized expression values.

### Transcription factor and gene regulatory network analysis

Transcription factor (TF) binding enrichment in open chromatin (ATAC) data was estimated using ChromVAR^47^ motif enrichment analysis with default parameters and cisBP-based position frequency matrices (PFM) from FigR^48^. Differential enrichment between groups of cells was calculated using Seurat’s *FindMarkers* function (Wilcoxon Rank Sum test with p-values adjusted using Bonferroni correction) on the ChromVAR deviation scores (Z scores).

The relationship between transcription factors and their target genes for building gene regulatory networks were inferred from both gene expression and ATAC data using FigR. FigR combines gene-peak correlation calculated from both modalities to identify domains of regulatory chromatin (DORCs) which are gene neighborhoods that likely have high regulatory activity. The algorithm then combines relative enrichment of TF motifs in DORCs, and the correlation of TF RNA expression and DORC accessibility to identify TFs that activate or repress different target genes. TFs that were most enriched in multiciliated and secretory lineages were selected based on ChromVAR enrichment scores and established literature on their role in FT cell lineages. Regulatory relationships between these TFs and their target genes inferred by FigR were summarized as gene regulatory networks using the ggnetwork R package.^86^

### Age-associated analyses of cell composition and organoid formation

Donor ages were obtained from clinical metadata. Ratios of epithelial to lineage-negative non-epithelial cells, multiciliated to secretory cells, and the percentage composition of individual epithelial clusters were compared with patient age using simple linear regression. Upon receipt, tissue samples were weighed, and following processing (as described above), total cell counts were obtained. Viable cells per milligram of tissue were calculated for each patient. In addition, the total number of organoids per well was recorded for each patient, enabling calculation of organoids per starting viable cell. Both viability-normalized organoid counts and viable cell counts per milligram of starting tissue were then correlated with patient age using simple linear regression.

### Secretome and proteomic profiling of FT tissue, 2D cultured cells, and 3D organoids

For secretome profiling, matched ampulla and fimbria tissues from two patients (four specimens total) were processed. Right ampulla and fimbria tissues were rinsed three times with PBS (Corning, 21-040-CV) (30 seconds each with gentle swirling), followed by three 30-minute washes in Advanced DMEM/F12 (Gibco, 1998031) at 37°C. Tissues were then transferred to fresh culture dishes and incubated in Advanced DMEM/F12 at 37°C for 24 hours. Conditioned media were subsequently collected, centrifuged at 200 × g for 10 minutes and 2,000 × g for 20 minutes to remove debris, and the resulting supernatant was snap frozen for downstream analysis. Following this, the tissue was transferred to a new culture dish and incubated in Advanced DMEM/F12 with 1X cOmplete Mini EDTA-free protease inhibitor cocktail (Roche Diagnostics, 11836170001) at 37°C for 24 hours. Conditioned media was collected as described above.

In parallel, left ampulla and fimbria tissues were dissociated as described above in FT specimen processing to obtain single-cell suspensions. Approximately one-quarter of the total cells were plated in FT epithelial cell culture medium and cultured to 60–70% confluence. Cultures were then washed three times with PBS, then incubated in Advanced DMEM/F12 at 37°C for 24 hours. Conditioned media was collected and processed as above. The remaining three-quarters of dissociated cells were used for epithelial cell enrichment. Cells were pelleted, stained with DAPI, CD45, CD31, EpCAM, and CD49f for 20 minutes at 4°C, and EpCAM⁺ cells were sorted using a BD FACSMelody cell sorter. Sorted cells were pelleted and snap frozen for proteomic analysis.

Proteins were precipitated from conditioned media by adding four volumes of cold acetone. After overnight incubation at -20 °C, protein pellets were reconstituted in 8 M urea (50 mM triethylammonium bicarbonate, pH 8.5) and subjected to reduction and alkylation using 10 mM dithiothreitol and 40 mM iodoacetamide, respectively. Enzymatic digestion was performed with the addition of Trypsin/Lys-C mix (Promega) followed by incubation at 37°C for overnight.

Around 100 ng of peptides were loaded onto Evotips (EV2011, Evotip Pure, Evosep) and analyzed using a timsTOF Ultra 2 mass spectrometer (Bruker Daltonics, Bremen, Germany) coupled to an Evosep ONE system (Evosep Biosystems). Peptides were separated using Whisper Zoom 20 SPD method (68-minute gradient) on an analytical column (15 cm x 75 µm, C18 1.7 µm, IonOpticks, AUR3-15075C18-CSI). Mass spectrometry data were acquired in DDA-PASEF mode with 10 PASEF scans per cycle. The mass range was set to 100-1700 m/z and ion mobility range was set to 0.7-1.3 V s cm^−2^. The ramp time was set to 100 ms with a 100% duty cycle. The raw mass spectrometry data were searched against the UniProt human protein database (20,461 entries) using MSFragger (version 4.0) embedded in the FragPipe suite (version 21.1). Two missed cleavages were allowed with a fully tryptic option and carbamidomethylation of cysteine was considered as a fixed modification and oxidation of methionine and acetylation of the protein N-terminal were set as variable modifications. PSM and protein validation were performed using PeptideProphet and ProteinProphet, respectively, at a 1% false discovery rate.

### Identification of epithelial subset-specific secreted molecules

A comprehensive reference list of human secreted proteins was compiled from multiple sources: (1) Human Protein Atlas (HPA) secretome datasets across diverse tissues, (2) reviewed UniProt entries annotated as secreted, (3) FT lavage proteome data,^87^ (4) 2D cultured PAX8⁺ epithelial cell secretome,^55^ (5) 2D cultured P0 FT epithelial cell secretome (6) fresh tissue secretome, (7) FT EpCAM+ cell proteome, (8) FT 3D organoid proteome and (9) a published FT proteome.^56^ Gene identifiers were harmonized across sources, and entries with ambiguous naming conventions were manually curated. These datasets were merged into a single non-redundant list of secreted molecules, with annotations specifying the number and type of sources supporting each gene.

To identify secreted molecules enriched in FT epithelial subpopulations, differential gene expression analysis was performed across epithelial subclusters using Seurat’s *FindAllMarkers* with the following parameters: min.pct = 0.1, and logfc.threshold = 0.1. The curated secretome gene list was intersected with differentially expressed genes from the epithelial subset clusters. Genes were annotated by the number of supporting datasets and ranked by cluster-specific log₂ fold change. The top and total secreted molecules per epithelial cluster were further evaluated.

Gene signatures representing the FT secretome and proteome were constructed by aggregating all proteins detected across each respective dataset. These aggregated gene sets were applied to the integrated epithelial subset using Seurat’s *AddModuleScore* function to calculate per-cell signature scores.

### TCGA HGSOC RNA-Seq correlation to FT epithelial clusters

RNA-seq data from The Cancer Genome Atlas Ovarian Cancer cohort (TCGA-OV) were downloaded via the TCGAbiolinks package.^88^ Raw STAR-aligned gene count matrices were prepared using *GDCprepare*, and Ensembl gene IDs were mapped to HGNC symbols using biomaRt.^89^ Gene-level expression values were subset to tumor samples with available molecular subtypes (differentiated, immunoreactive, mesenchymal, and proliferative), which were obtained from UCSC Xena and linked to TCGA sample barcodes. Count matrices were normalized using edgeR,^90^ and expression values for the top secreted cluster markers were z-score transformed. The top 20 secreted molecules per epithelial subcluster were prioritized based on differential expression rank and retained if detected in TCGA samples. Expression profiles were visualized using ComplexHeatmap,^91^ grouped by TCGA molecular subtype and hierarchically clustered by epithelial subcluster.

### CPTAC HGSOC proteome correlation to FT epithelial clusters

Proteomic data from the CPTAC ovarian cancer cohort was downloaded.^62^ Normalized protein abundance values were obtained from the published matrix and filtered to remove non-gene entries and duplicated sample identifiers. Protein expression data were matched to TCGA sample barcodes with subtype annotations. The top 20 differentially expressed secreted molecules per epithelial cluster—excluding ribosomal and housekeeping genes—were used for analysis. Genes not detected in the CPTAC dataset were excluded, and the final matrix was Z-score normalized across samples. Expression profiles were visualized using ComplexHeatmap,^91^ grouped by TCGA molecular subtype and hierarchically clustered by epithelial subcluster.

### Comparison of FT epithelial secreted molecules with ovarian cancer tumor regions via microarray expression

To assess the expression of secreted epithelial molecules in laser capture microdissected (LCM) tumor regions, publicly available Affymetrix microarray data from Tothill et al^63^ was analyzed. The series matrix file was downloaded from the Gene Expression Omnibus (GEO) using the GEOquery R package.^92^ Probe-level data were mapped to HGNC gene symbols using the hgu133plus2.db Bioconductor annotation package.

A subset of 10 tumor samples consisting of paired epithelial and stromal LCM regions was used. Expression matrices were subset accordingly, and a gene signature representing the top 50 secreted molecules from epithelial cluster 5, identified via differential gene expression in scRNA-seq data, was applied. Genes not detected in the microarray platform were excluded, and the final matrix was Z-score normalized across samples. Heatmaps were generated using the pheatmap^93^ package in R, displaying scaled expression of signature genes across all selected samples.

### Comparative transcriptomic profiling of FT epithelial cells and fetal mesonephros

ScRNA-seq data from second-trimester female mesonephros were processed using Seurat.^65^ Raw 10X HDF5 files were imported with Read10X_h5, and cells were filtered to retain those with 200–8,000 genes, <20% mitochondrial transcripts, >1% ribosomal content, and <0.5% hemoglobin content. Normalization, variable gene selection, scaling, PCA, clustering (resolution=0.5), and UMAP were performed using standard Seurat workflows described above. Clusters were annotated using canonical markers identified by Taelman et al. To assess similarity between fetal and adult FT cell states, we computed module scores using *AddModuleScore*. The top 50 marker genes from FT cluster 5 (ranked by log2FC) were scored across fetal mesonephros cells.

## Acknowledgements

We thank study coordinators Katie Reed, Stephanie Hafner, Kaitlyn Schwartz, Susan Holtegaard, Autumn Moon, and Emily Bruetzman for their invaluable support, and Trace Christensen for his technical assistance with electron microscopy. We thank Ishaq Viringipurampeer for his technical assistance. We are also incredibly grateful to our patients and surgical colleagues for contributing tissue samples to the biobank.

