## Supplementary figures and images for "A Rare Multipotent Peg-like Epithelial Cell is a Candidate Cell-of-Origin for High-Grade Serous Ovarian Cancer"

### Supplementary Figure 1

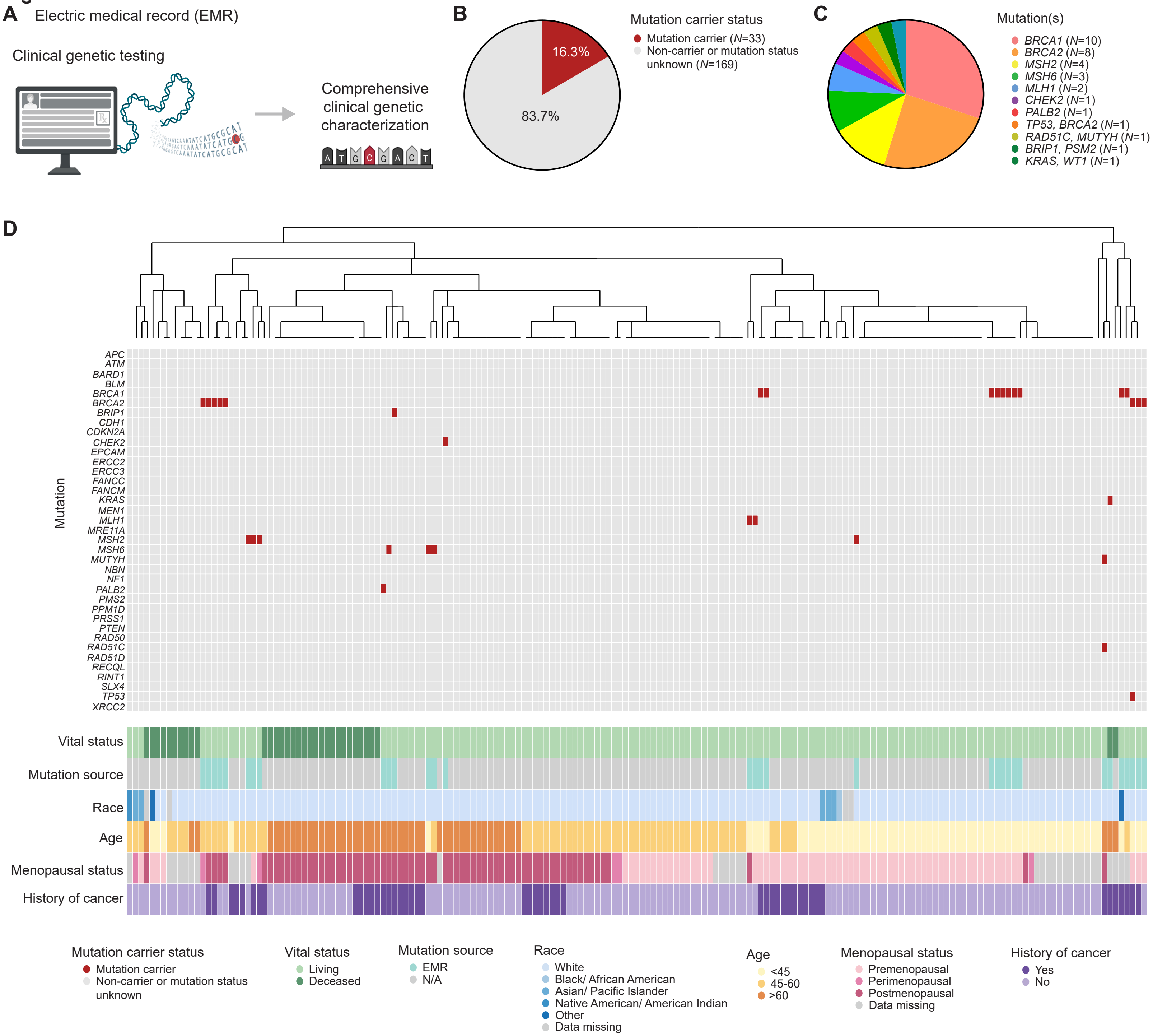

### Supplementary Figure 2

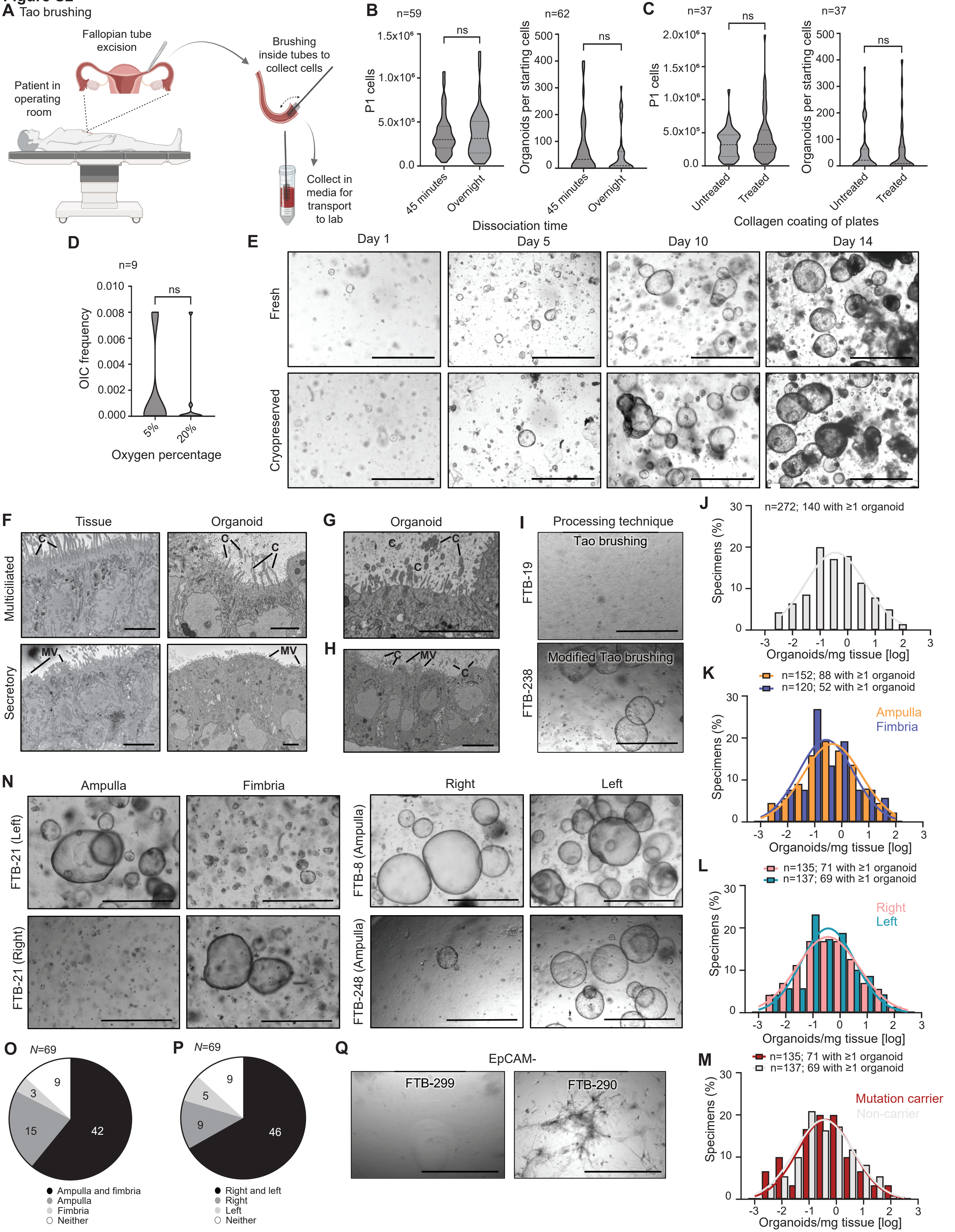

### Supplementary Figure 3

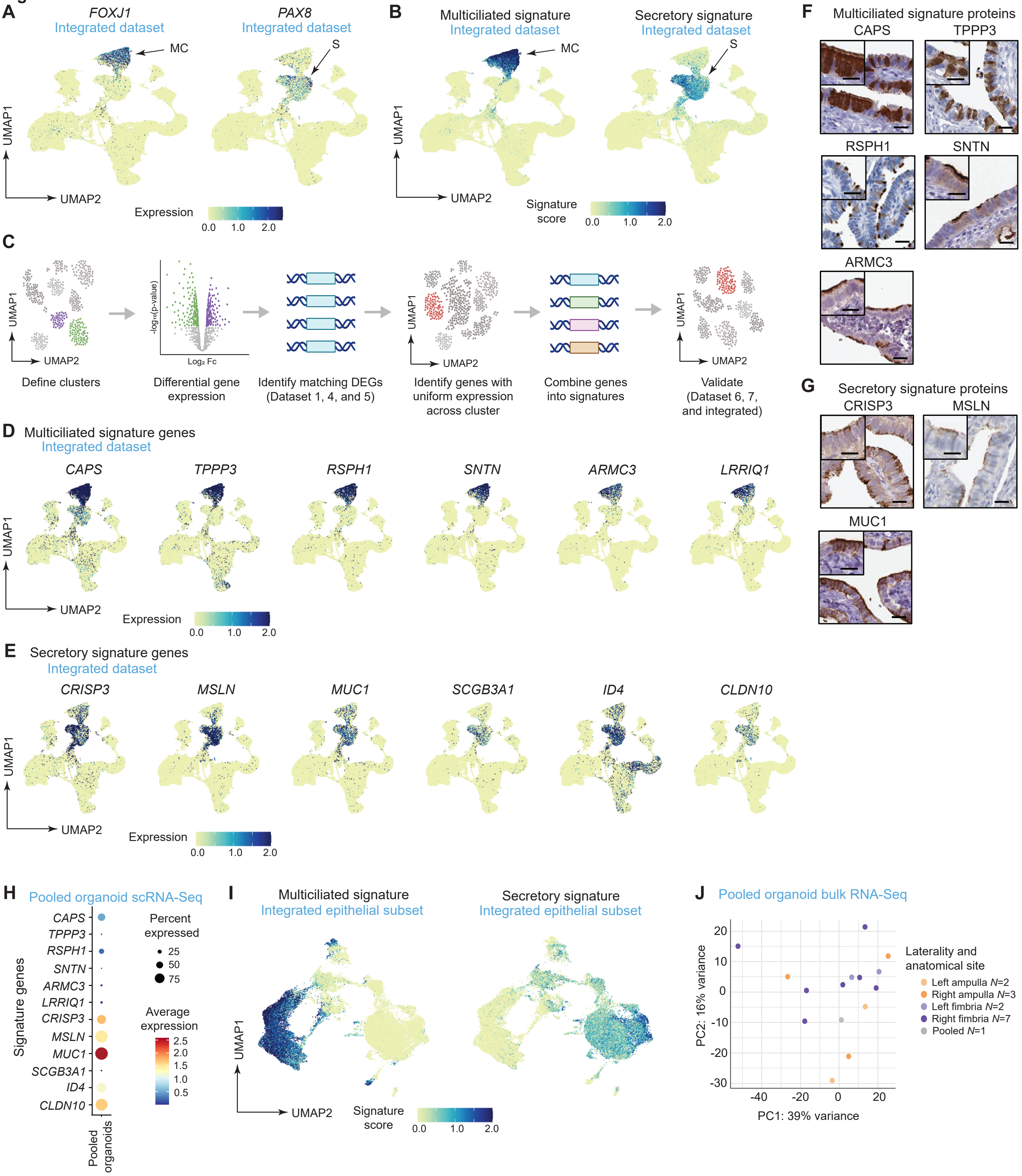

### Supplementary Figure 4

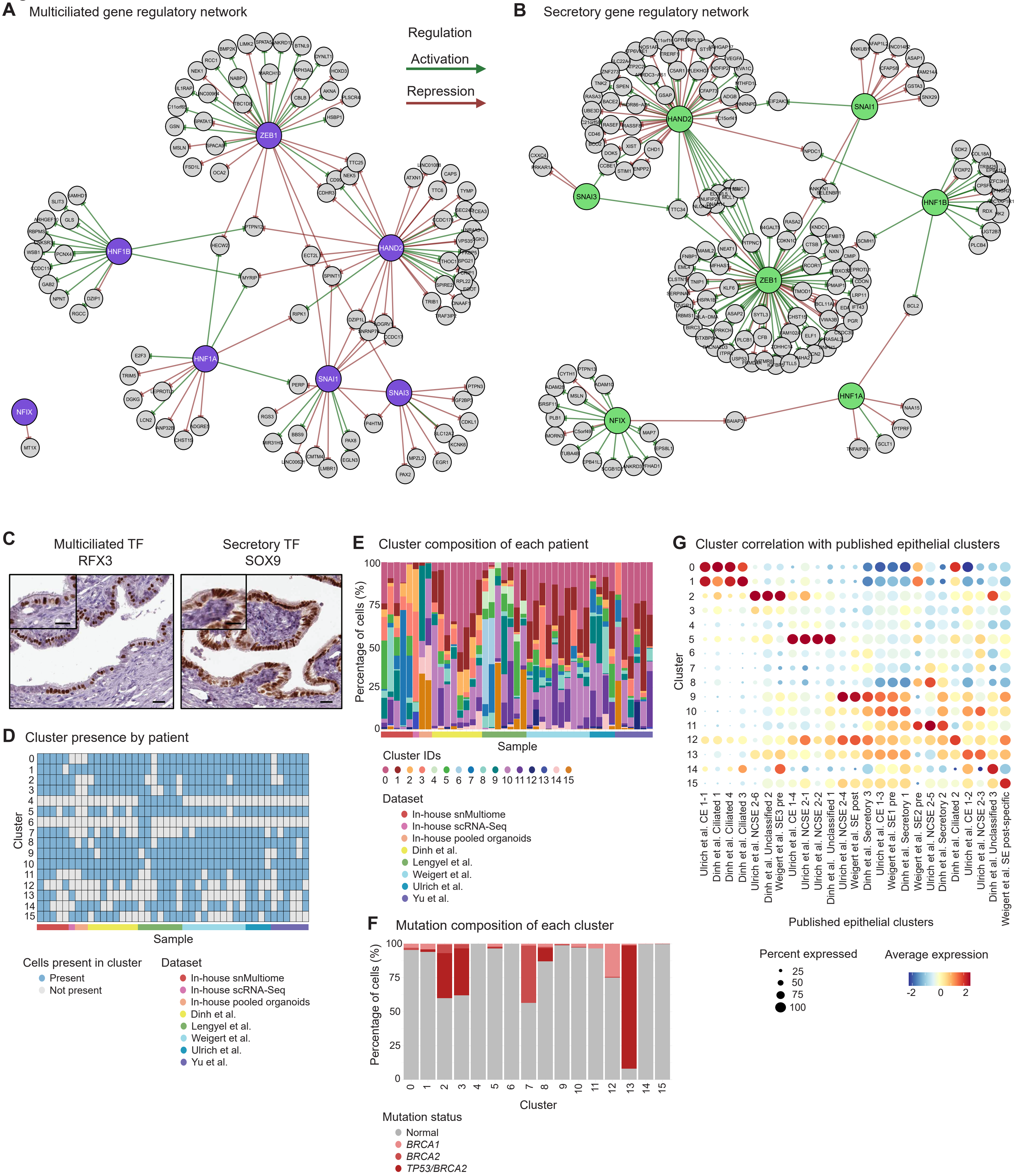

### Supplementary Figure 5

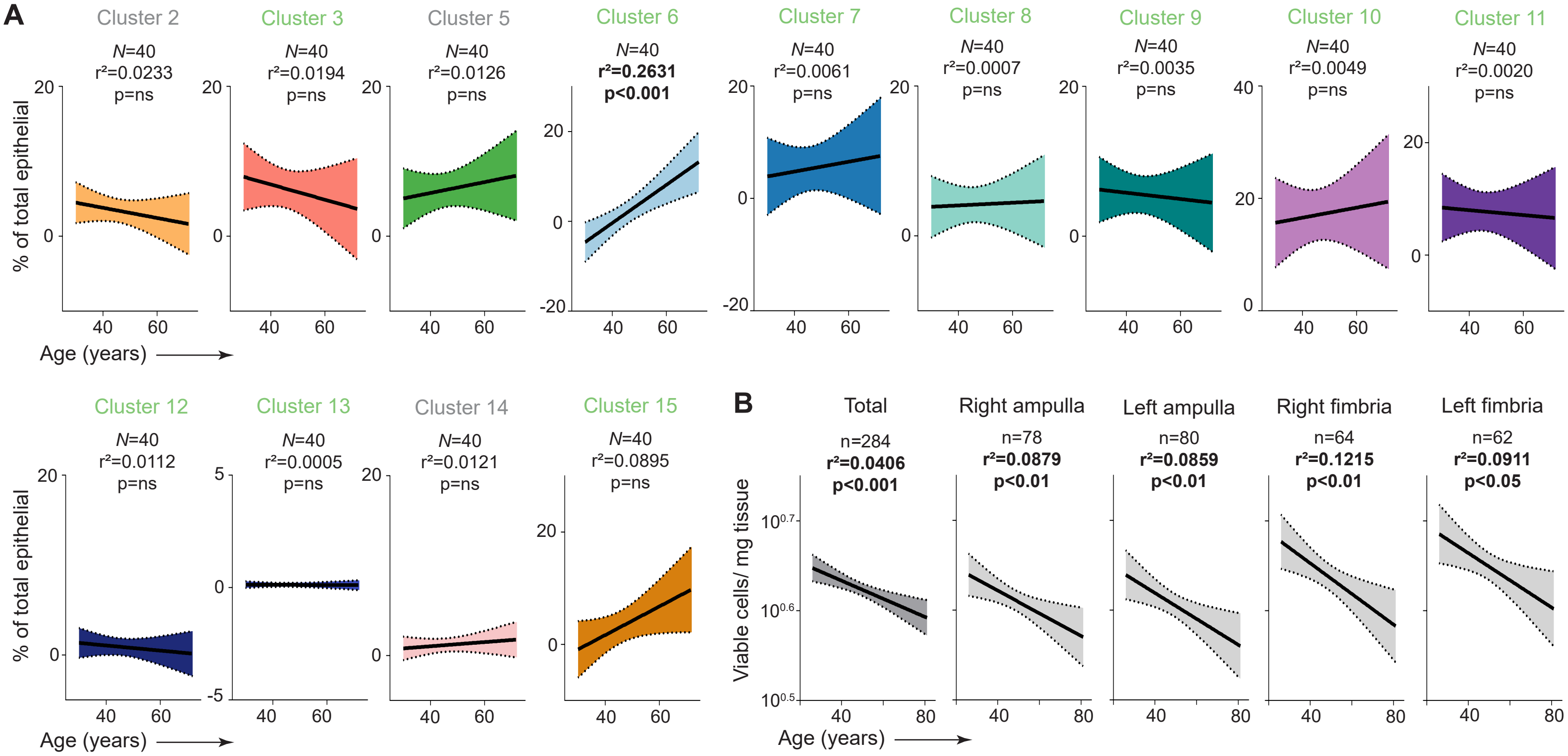

### Supplementary Figure 6

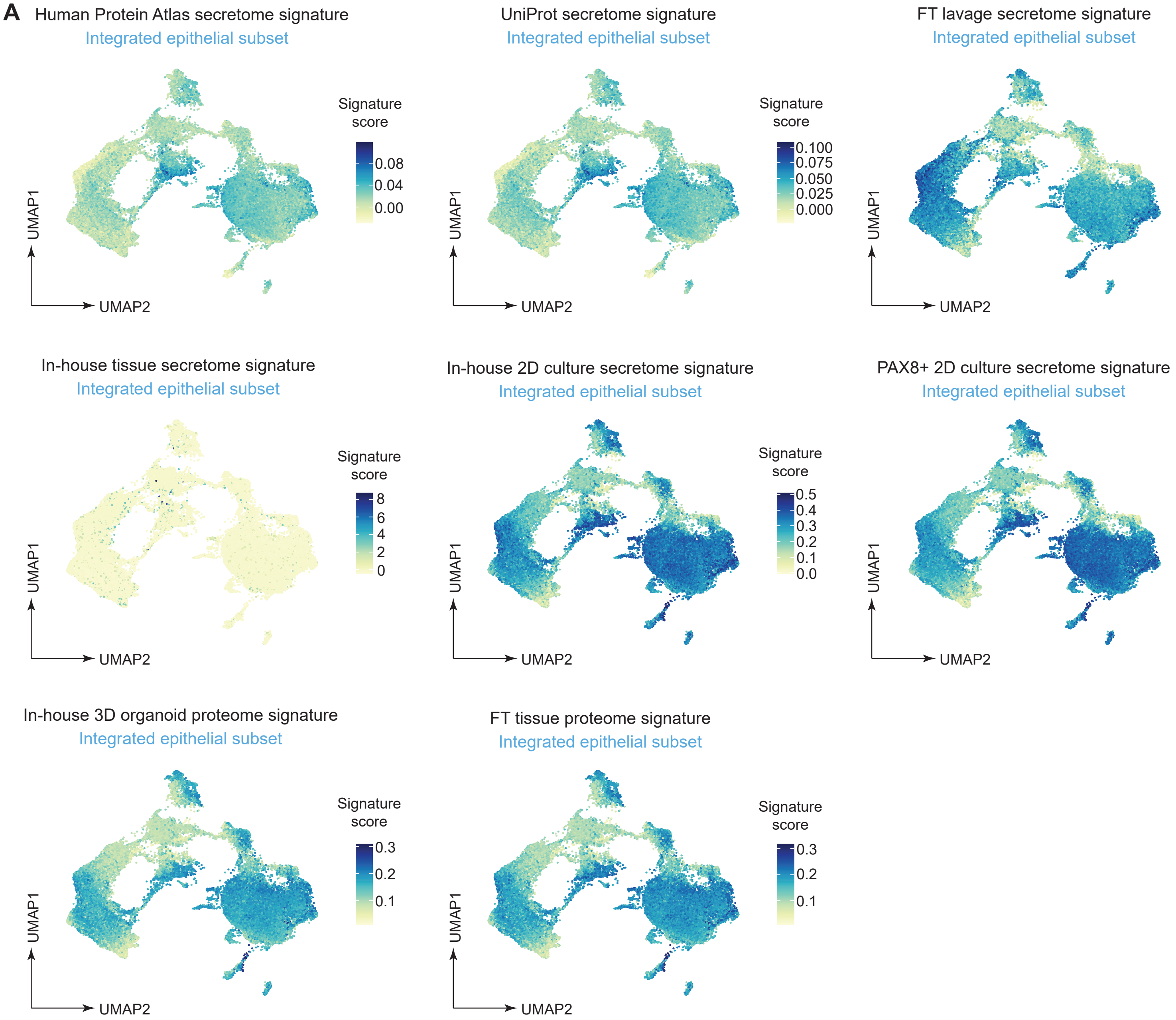

### Supplementary Figure 7

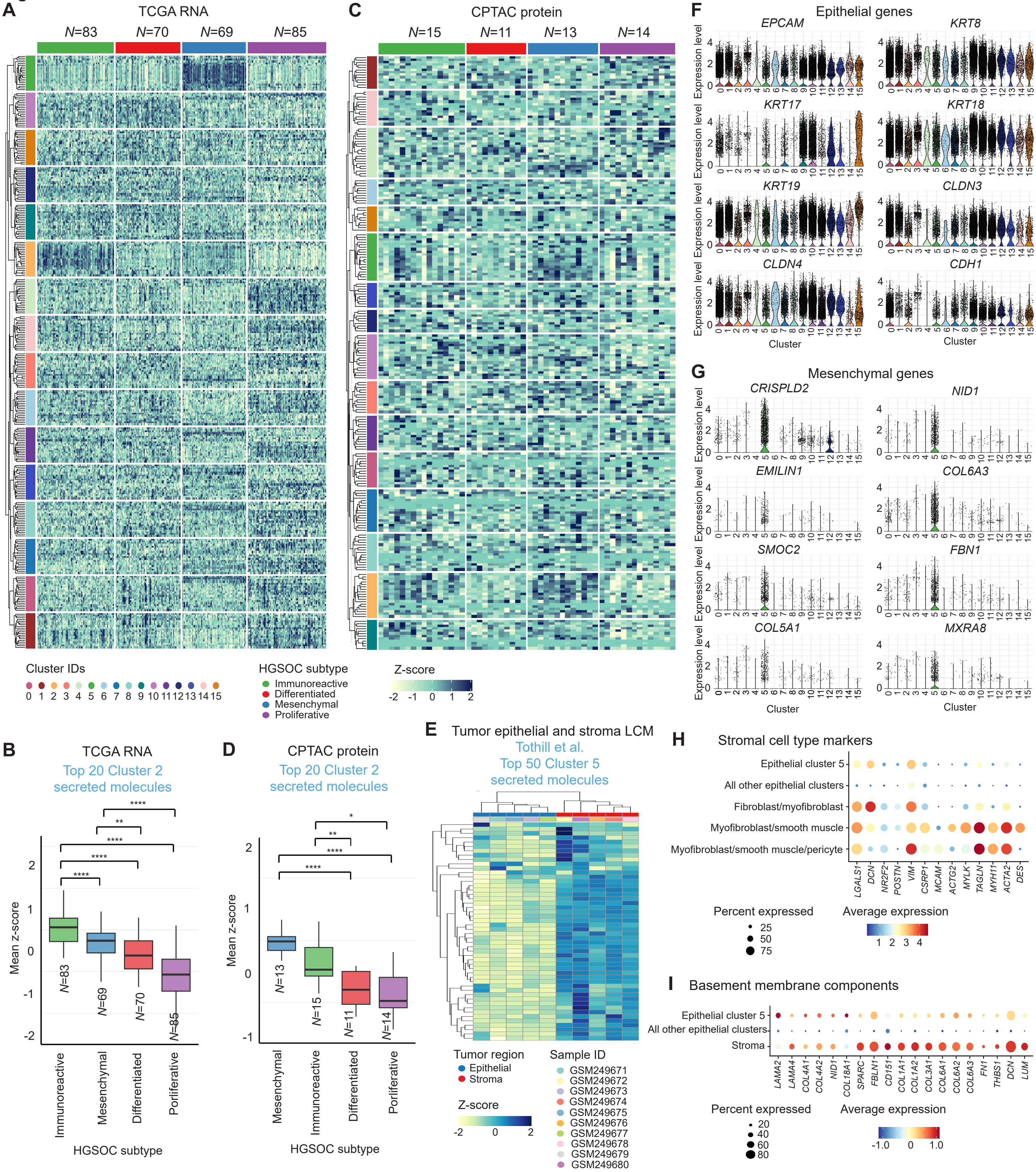
